# Suppressive might of a few: T follicular regulatory cells impede auto-reactivity despite being outnumbered in the germinal centres

**DOI:** 10.1101/2023.07.10.548342

**Authors:** Marta Schips, Tanmay Mitra, Arnab Bandyopadhyay, Michael Meyer-Hermann

## Abstract

The selection of high-affinity B cells and the production of high-affinity antibodies are mediated by T follicular helper cells (Tfhs) within germinal centres (GCs). Therein, somatic hypermutation and selection enhance B cell affinity but risk the emergence of self-reactive B cell clones. Despite being outnumbered compared to their helper counterpart, the ablation of T follicular regulatory cells (Tfrs) results in enhanced dissemination of self-reactive antibody-secreting cells (ASCs). The specific mechanisms by which Tfrs exert their regulatory action on self-reactive B cells are largely unknown. We developed computer simulations to investigate how Tfrs regulate either selection or differentiation of B cells to prevent auto-reactivity. We observed that Tfr-induced apoptosis of self-reactive B cells during the selection phase impedes self-reactivity with physiological Tfr numbers, especially when Tfrs can access centrocyte-enriched GC areas. While this aided in selecting non-self-reactive B cells by restraining competition, higher Tfr numbers distracted non-self-reactive B cells from receiving survival signals from Tfhs. Thus, the location and number of Tfrs must be regulated to circumvent such *Tfr distraction* and avoid disrupting GC evolution. In contrast, when Tfrs regulate differentiation of selected centrocytes by promoting recycling to the dark zone phenotype of self-reactive GC resident pre-plasma cells (GCPCs), higher Tfr numbers were required to impede the circulation of self-reactive ASCs (s–ASCs). On the other hand, Tfr-engagement with GCPCs and subsequent apoptosis of s–ASCs can control self-reactivity with low Tfr numbers, but does not confer selection advantage to non-self-reactive B cells. The simulations predict that to restrict auto-reactivity, natural redemption of self-reactive B cells is insufficient and that Tfrs should increase the mutation probability of self-reactive B cells.

## 1 Introduction

To mount an effective immune response, it is crucial to produce antibodies that specifically target foreign antigens while avoiding the production of self-reactive antibodies. This delicate balance is achieved in a specialized micro-environment within secondary lymphoid organs, referred to as germinal centres (GCs). Within the GC, B cells undergo affinity maturation, an evolutionary process which ensures the generation of high-affinity and specific antibodies. Affinity maturation is achieved by cycles of somatic hypermutation (SHM) of the B cell receptor (BCR) and subsequent selection based on its affinity. This selection process heavily relies on the interactions of B cells with T follicular helper cells (Tfhs), which promote the expansion of B cell clones with the best-fit receptor. The outcome of the GC reaction encompasses the generation of memory B cells, capable of rapid reactivation upon secondary exposure, and plasma cells (PCs or antibody secreting cells (ASCs)) that secrete antibodies and home to the bone marrow to employ long-term protection.

Apart from Tfhs, emerging studies have highlighted the pivotal role of T follicular regulatory cells (Tfrs) in the GC micro-environment. Tfrs differentiate from CD25^hi^Foxp3^+^ regulatory T cells (Treg) [1, 2], but share signalling pathways involved in Tfh cell development [1, 3]. Notably, Tfrs can be distinguished based on their CD25 expression, with CD25^+^ Tfrs predominantly located at the GC border and CD25^*−*^ Tfrs primarily found inside the GC [2, 4]. Tfrs exhibit a phenotype which resembles Tfhs, expressing programmed cell death protein-1 (PD-1), C-X-C chemokine receptor type 5 (CXCR5), and B cell lymphoma 6 (Bcl6) to name a few [1, 3, 5]. However, unlike Tfhs, Tfrs lack co-stimulatory molecules, such as IL-21 and IL-4 [1, 3, 5, 6]. Instead, they express Treg-specific molecules, including cytotoxic T lymphocyte-associated protein 4 (CTLA-4), and Foxp3 [1, 4]. Functionally, Tfrs exert suppressive effects on the GC micro-environment to limit the expansion of self-reactive antibodies [1, 2, 3, 5, 7, 8, 9, 10].

The critical role of Tregs in maintaining self-tolerance has long been established [11]. Analogously, Tfrs have emerged as essential regulators of self-tolerance within the GC [12, 13, 14, 15]. The process of SHM, which introduces point mutations into the variable regions of BCR immunoglobulin genes, is essential for enhancing B cell affinity but also carries the risk of generating self-reactive BCRs [16, 17]. Studies in mice lacking Tfrs during influenza-virus infection and protein immunization have demonstrated the development of s–ASCs and the circulation of auto-antibodies such as IgG and IgE [2, 18]. Furthermore, Tfr-deficient mice have been shown to develop auto-antibodies at an older age [8, 19]. Although studies have implicated CTLA-4 [6, 20, 21], neuritin [19] and IL-10 [22] as potential mediators of Tfr functions, the specific mechanisms by which Tfrs exert their regulatory action on self-reactive GC B cells (s–GCBCs) are largely unknown. Moreover, the effects of Tfr depletion on antibody responses in different experimental settings have yielded conflicting results, with some studies reporting enhanced antigen-specific antibody responses in the absence of Tfrs [3, 5] and others showing the opposite effect [1, 9, 18, 19].

The precise role of Tfrs within and outside the GC remains unclear. Recent research has challenged the notion that CXCR5 is necessary for accessing the GC, and reported that reducing the number of Tfrs did not significantly impact the GC output [23]. Intriguingly, experimental studies have suggested that Tfrs may be actively attracted into the GC by chemokines such as CCL3 [24]. Furthermore, a subset of Tfrs appears to emerge as a consequence of Tfh differentiation at a later stage of the GC reaction [25]. Additionally, the question of how a relatively small population of Tfrs can exert multiple and sometimes discordant effects remains unanswered, particularly considering the variable ratios of Tfhs to Tfrs reported in the literature.

To address these complexities, we have developed an agent-based model that explores different checkpoints within the GC reaction at which Tfrs may regulate self-reactivity arising from SHM. Our computational study takes into account the relative proportions of Tfhs to Tfrs and their spatial localization within the GC. Our findings indicate that a direct intervention by a moderate number of Tfrs in the selection process of s–GCBCs can prevent auto-reactivity in physiological Tfr numbers. While this may aid affinity maturation of non-self-reactive GC B cells (ns–GCBCs) by restraining competition, a higher number of Tfrs can engage with ns–GCBCs and result in *Tfr distraction* by hindering them to receive survival signals from Tfhs, thereby attenuating selection of ns–GCBCs. Thus, the number or location of Tfrs must be regulated to avoid disrupting the selection process of ns–GCBCs and GC evolution. Moreover, the localization of a low number of Tfrs in the GC areas enriched in centrocytes (CCs), especially when more s–GCBCs are present (e.g., during the expansion phase of the GC), seems crucial to prevent auto-reactivity while facilitating affinity maturation of ns– GCBCs. We further propose that Tfrs can also impede the circulation of s–ASCs by acting at the GC B cell differentiation step during the GC response. A higher number of Tfrs was necessary to control s– ASCs by promoting dark zone (DZ) recycling of self-reactive germinal centre resident pre-plasma cells (GCPCs), and resulted in reduced affinity and numbers of ns–ASCs. Conversely, engagement of Tfrs with GCPCs and subsequent apoptosis of s–ASCs could effectively control self-reactivity even with a lower number of Tfrs, but did not confer any advantage to ns–GCBCs during selection. Understanding the precise mechanisms by which Tfrs regulate self-reactivity within the GC is of paramount importance for deciphering the intricate interplay between B and T cell interactions during the immune response. This knowledge holds promise for advancing our understanding of autoimmune diseases and optimizing vaccine strategies. Furthermore, unravelling the complex role of Tfrs in the GC reaction will provide valuable insights into immune regulation and tolerance mechanisms.

## 2 Results

Building upon our prior investigations [26, 27, 28], we developed an agent-based model of the GC reaction, which, to the best of our knowledge, is the first to study how Tfrs suppress the emergence of auto-reactivity arising from the s–GCBCs clones generated through SHM.

### 2.1 Redemption is insufficient to control self-reactivity in GC

In our *in silico* framework, B cells divide in the DZ and can mutate at each division with an affinity-dependent probability (Fig. 1A and Methods). We consider each mutation to be associated with a constant probability (*pSelf*) of generating a s–GCBC. As suggested by experiments [29], we introduced a probability (*pRed*) for s–GCBCs to re-edit their BCRs following a mutation in the DZ and lose self-reactivity. Unless otherwise stated, we regarded *pSelf* =*pRed*. The framework assumes that the daughter cells of s–GCBCs inherit self-reactivity if unredeemed.

**Figure 1:**
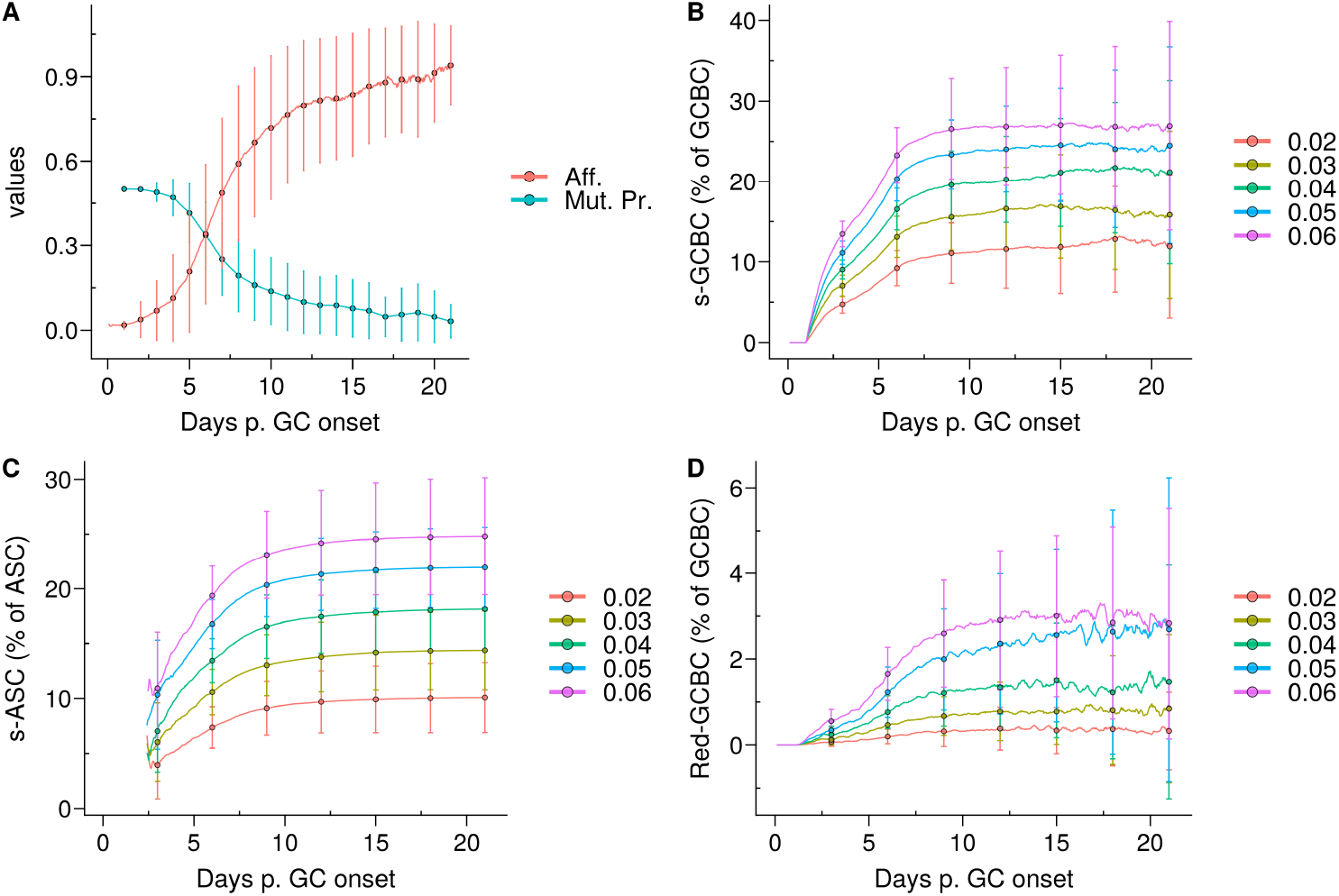
Emergence of self-reactivity in the GC. **(A)** Evolution of affinity (Aff.) of live germinal centre B cells (GCBCs) (red) and mutation probability (Mut. Pr.) of dark zone B cells (cyan); **(B)** Kinetics of the quantity of self-reactive GCBCs (s–GCBCs) as a percentage of total GCBCs; **(C)** Kinetics of the quantity of self-reactive antibody secreting cells (s–ASCs) as a percentage of total ASCs; **(D)** Kinetics of the quantity of redeemed-GCBCs as a percentage of total GCBCs. In each panel, the lines and points denote the mean, whereas the error bars depict the standard deviations of 96 simulations; in **(B–D)**, each colour corresponds to a different probability *pSelf* to generate s–GCBCs during somatic hypermutation.

CCs interact with FDCs to collect antigens and with Tfhs to receive survival signals in the light zone (LZ). In our study, we operated under the assumption that B cells undergo affinity maturation regardless of their self-specificity. Consequently, in the absence of any regulatory mechanism, s–GCBCs compete with ns–GCBCs for selection and eventually differentiate into self-reactive antibody-secreting cells (s–ASCs) post selection.

The emergence of self-reactive clones in the GC correlated with the mutation probability, resulting in an increase in the self-reactive population until ∼day 9, ranging from 10% to 25% depending on the *pSelf* value. Afterwards, the self-reactive population remained relatively stable due to the marginal mutation probability (Fig.1B). The percentage of s–ASCs mirrored the dynamics of s–GCBCs (Fig.1C). Similarly, the population of redeemed B cells that rescued their specificity for non-self, increased alongside the s–GCBC population, but it remained a minor fraction of the overall GC population (Fig. 1D).

Therefore, under the aforementioned assumptions, the model suggests that the majority of s– GCBCs is generated early during the GC reaction and that a natural redemption process is not sufficient to prevent the circulation of s–ASCs. This finding underscores the necessity for additional extrinsic or distinct mechanisms to impede dissemination of s–ASCs.

### 2.2 Tfr mediated regulation in GCs

Experimental results consistently demonstrated that depletion of Tfrs led to enhanced circulation of self-reactive antibodies and induced autoimmunity in mice [1, 2, 8, 10, 18, 19]. Therefore, we addressed how Tfrs could interfere with the selection process of s–GCBCs and prevent spreading of self-reactive antibodies.

CCs acquire antigens from FDCs and subsequently receive survival signals from Tfhs crucial for their selection. Upon selection, each B cell may upregulate factors associated with final differentiation to GC plasma cells (GCPCs), e.g., CD138. GCPCs differentiate into ASCs after ∼12 hours and exit the GC. On the other hand, CD138^*−*^ CCs recycle back to the DZ phenotype after ∼6 hours, preparing for another round of proliferation and affinity maturation (see Methods).

We developed three different models of Tfrs acting on distinct stages of the selection or differentiation process (see Methods):

- *Apoptosis* model (Fig. 2B): Tfrs primarily originate from natural Tregs [1, 3, 5]. Consequently, their T cell receptor repertoires closely resemble those of natural Tregs [30], indicating a predominant presence of self-specificity. Accordingly, we hypothesized that Tfrs survey the GC micro-environment to suppress s–GCBCs by interacting with s–GCBCs that successfully acquired antigens and are searching for Tfhs to receive survival signals. In this model, engagement with Tfrs is assumed to lead to apoptosis specifically in s–GCBCs, while being innocuous to ns–GCBCs.

**Figure 2:**
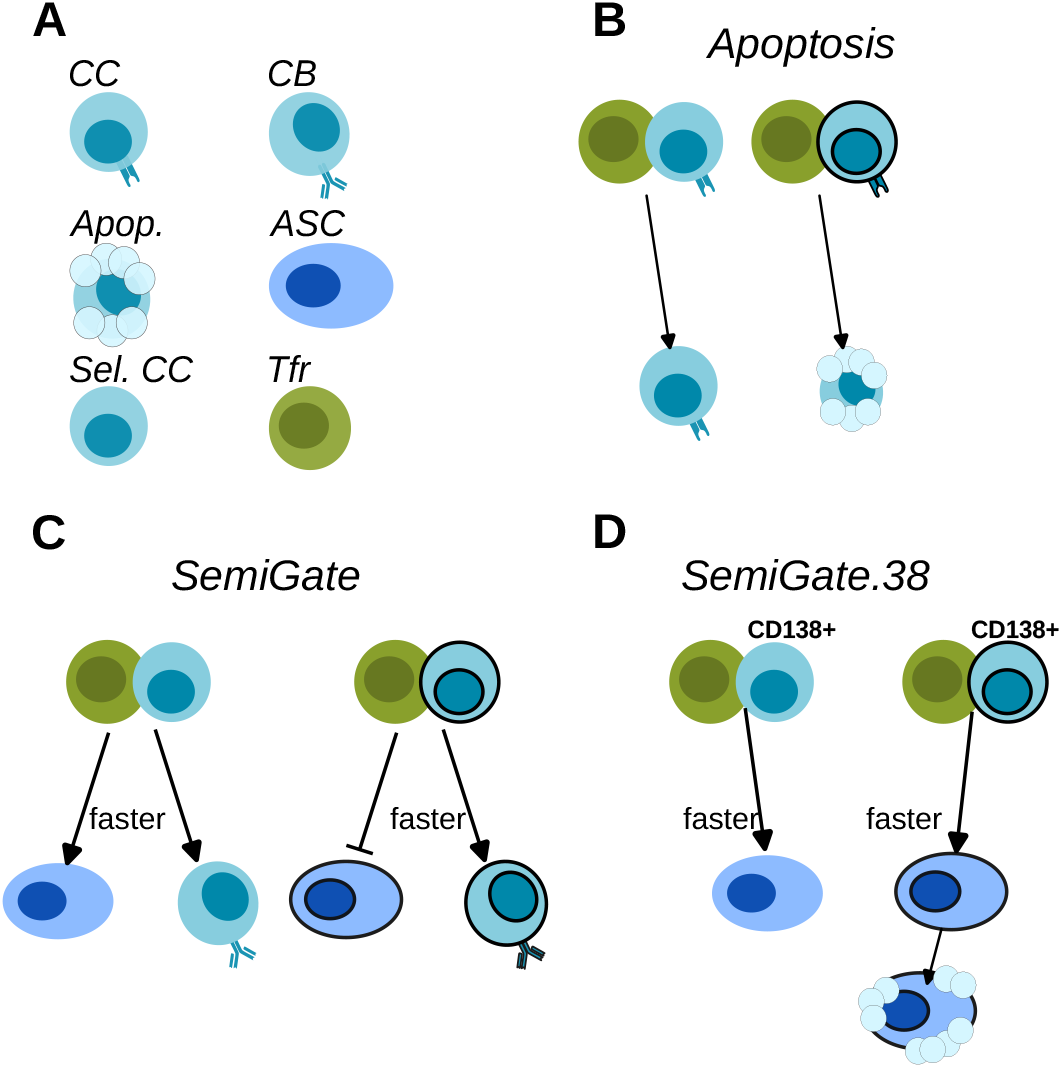
Schematics of Tfr action on the B cells. **(A)** Cell-type legend: MHC-expressing BCs are FDC-selected CCs (CC), BCR-expressing BCs are CBs (CB), Apoptotic cells (Apop.), antibody-secreting-cells (ASC), Tfh-selected CCs (Sel. CC), T follicular regulatory cells (Tfr); **(B)** *Apoptosis* model; **(C)** *SemiGate* model; **(D)** *SemiGate*.*38* model. In **(B–D)** black-border indicates self-specificity for each type of cells. In **(C–D)** thicker lines indicate a faster differentiation following Tfr interaction.

In recent Tfr-depletion experiments, a significant accumulation of a distinct population of GCBCs with a plasma cell (PC) phenotype, referred to as GCPCs, was observed. The GCs lacking Tfrs exhibited approximately seven times higher numbers of GCPCs compared to the control group [19]. These GCPCs expressed CD138 and showed elevated levels of Blimp-1, along with decreased levels of Bcl6, while maintaining expression of B220. Despite the increased presence of GCPCs, the overall number of PCs did not decrease on either day 7 (post SRBCs immunization) or day 16 (post-NP-OVA immunization) when Tfrs were depleted [19]. Moreover, the evidence demonstrated that Tfrs preferentially interact with CCs that secrete CCL3 [24]. Considering that CCL3 secretion is repressed by Bcl6 [31], we postulated that Tfh-selected CCs, which downregulate Bcl6, would attract Tfrs for interaction. These findings led us to consider the possibility that Tfrs may exert their regulatory effects following Tfh mediated selection, when the fate of B cells regarding DZ recycling versus ASC differentiation is determined. We explored this hypothesis by implementing the *SemiGate* and *SemiGate*.*38* models in which Tfrs interact exclusively with Tfh-selected CCs. The main differences between the two models are described below.

- *SemiGate* model (Fig. 2C): In the *SemiGate* model, the interaction between Tfrs and Tfh-selected CCs yields distinct outcomes depending on CD138 expression and specificity for self. ns–GCPCs undergo prompt differentiation into ASCs in response to interaction with Tfrs. Conversely, s– GCPCs and CD138^*−*^ CCs are directed to recycle back to the DZ phenotype.
- *SemiGate*.*38* model (Fig. 2D): A few experimental studies suggested that auto-reactivity may not be censored within the GC, and revealed an apoptosis-dependent checkpoint targeting auto-reactive memory and plasma cells outside the GC environment [32, 33]. Motivated by these findings, we investigated the potential impact of Tfr interactions on GC output. In this model, Tfrs selectively engage with GCPCs, facilitating their differentiation into ASCs. Furthermore, Tfr interactions induce apoptosis in s–ASCs after the GC stage. This notion finds further support from *in vitro* studies demonstrating altered B cell metabolism and impaired antibody secretion upon Tfr interactions [9].

#### 2.2.1 Motility and localization of Tfrs

Tfrs are identified as CXCR5^+^FoxP3^+^PD1^+^CD4^+^ T cells. While Tfrs located at the T-zone:B-zone border around the GC are found to be predominantly CD25^+^, to access the GC, Tfrs downregulate CD25 [4]. Generally, the experimental studies involving Tfr-depletion or genetic manipulation do not distinguish Tfrs based on their CD25 expression. Therefore, it is unclear how the localization and motility of Tfrs impact the GC evolution and exert protection against auto-reactivity. To investigate this, we considered two types of motility of Tfrs as described below (see Methods).

- *Tfh-like* motility: Resembling motility of Tfhs, Tfrs move randomly in the GC with a preferential directionality for the LZ edge (Supplementary Material, Fig. S1A, left subpanel);
- *CC-like* motility: Resembling motility of CCs [34], Tfrs re- and de-sensitise for CXCL13 in a concentration-dependent manner (Supplementary Material, Fig. S1A, right subpanel);

Depending on their motility, Tfrs resolve in distinct spatial distributions (Supplementary Material, Fig. S1B,C), which result in different degrees of Tfr:GCBC engagement (Fig. 3).

**Figure 3:**
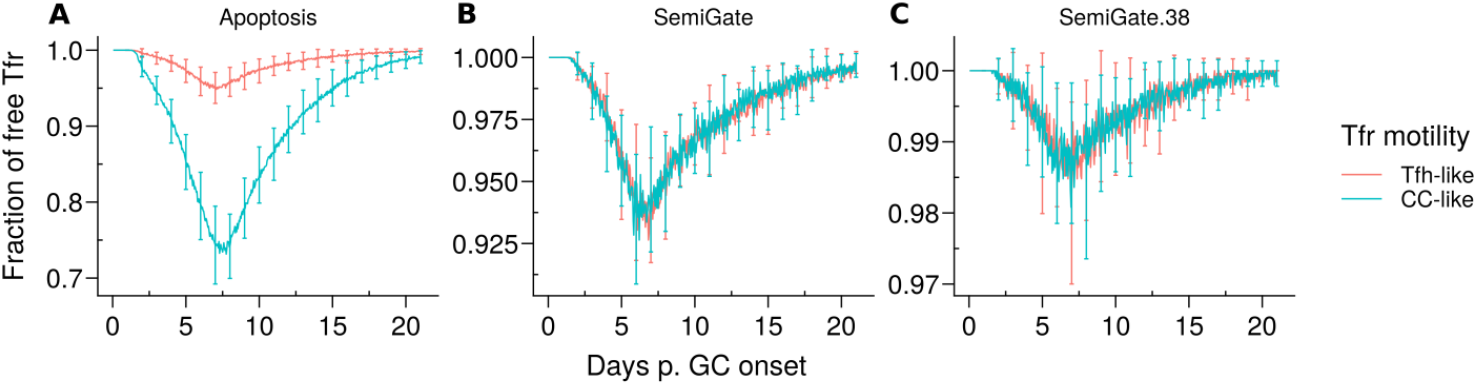
Representative dynamics of Tfr engagement under different motility types and Tfr regulation models. **(A)** *Apoptosis* model, in which Tfrs interact with germinal centre B cells (GCBCs) during selection; **(B)** *SemiGate* model, in which Tfrs interact only with selected centrocytes (CCs); **(C)** *SemiGate*.*38* model, in which Tfrs interact only with germinal centre resident pre-plasma cells (GCPCs). Colours represent different motility types of Tfrs: *Tfh-like* (red), *CC-like* (cyan). Lines and error bars represent, respectively, mean and standard deviation of 96 simulations. The results were obtained with *pSelf* = 0.04 and Tfh:Tfr = 2:1.

It is noteworthy that, in the *Apoptosis* model, the proportion of unbound Tfrs was significantly lower when Tfrs had access to the CC-enriched GC area (Fig. 3A, *CC-like* motility). This was attributed to the co-localization of Tfrs and CCs (Supplementary Material, Fig. S1B, right panel). In contrast, in other models, Tfrs interact only with those B cells which were positively selected by Tfhs and start moving randomly in the GC before differentiating in the DZ or output phenotype. Hence, the interaction opportunities between Tfrs and B cells are comparable in the scenarios of *Tfh-like* and *CC-like* motility. These observations imply that the spatial organization of Tfrs is likely to have a significant impact on the outcomes of the *Apoptosis* model.

### 2.3 Tfr-induced s–GCBC apoptosis suppresses auto-reactivity but may disrupt ns–GCBC selection

Using the *Apoptosis* model, we investigated how induction of apoptosis in s–GCBCs following their interactions with Tfrs impact the dynamics and self-reactivity in the GC.

When Tfrs were spread in the LZ (*Tfh–like* motility), the GC volume remained comparable to the scenario without Tfrs (Tfh:Tfr - 250:0) (Fig. 4A, left panel). However, when Tfrs resembled the motility of the CCs, the progression of the GC was compromised at higher Tfr numbers (corresponding to lower Tfh:Tfr ratios) (Fig. 4A, right panel). In both scenarios, the frequency of the selected s–GCBCs significantly reduced with increasing numbers of Tfrs (Supplementary Material, Fig. S2A).

**Figure 4:**
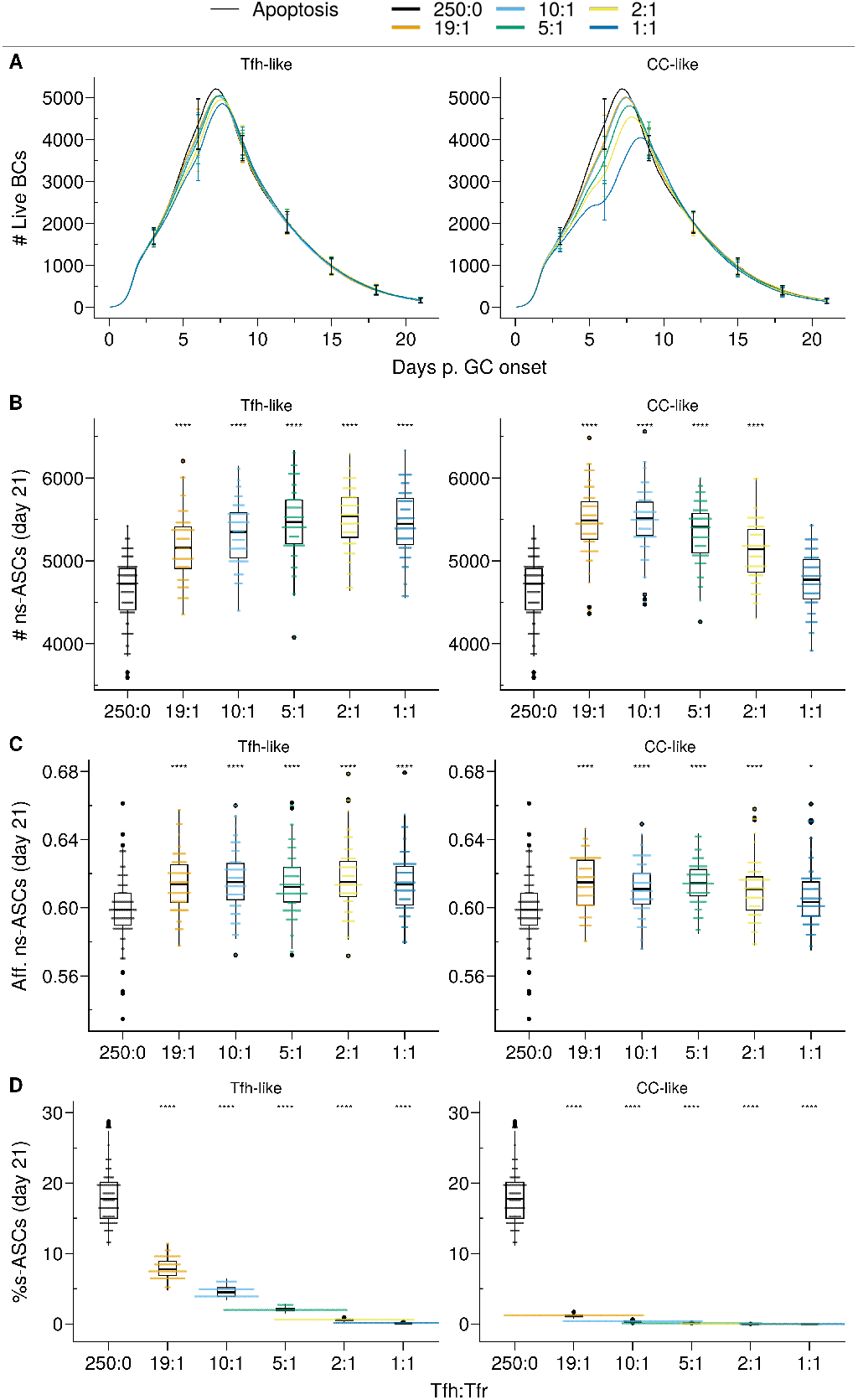
*Apoptosis* model suppresses auto-reactivity at physiological Tfr numbers. **(A)** Germinal centre B cell (GCBC) kinetics; **(B)** Number of non-self-reactive antibody secreting cells (ns–ASCs); **(C)** Affinity of ns–ASCs; **(D)** Percentage of self-reactive ASCs (s–ASCs) in total ASCs. In **(B–D)** the analysis was performed at day 21 post-GC onset. Each column shows the results for different motility types: *Tfh-like* (left coloumn), *CC-like* (right coloumn). Each colour depicts a different Tfh:Tfr ratio. In **(A)**, lines and error bars are means and standard deviations, respectively. SD at days 3,6,9,12,15,21. In **(B–D)**, each dot represents a different simulation, and significance is denoted with Wilcoxon signed rank tests against the group with no Tfr (Tfh:Tfr ratio = 250:0). Statistics were performed on 96 simulations. The results were obtained with *pSelf* =0.04.

In the *Tfh-like* motility, Tfrs promoted the selection and expansion of ns–ASCs, resulting in increased production of ns–ASCs with enhanced affinity (Fig. 4B,C right panel). We observed a similar trend when Tfrs obtained preferential access to the area rich in CCs relative to Tfhs (*CC-like* motility). In this setting, at a moderate number of Tfrs, the expansion of ns–ASCs was prominent and corresponded to enhanced affinity (Fig. 4B,C, left panel). Conversely, at a higher number of Tfrs, the increased interaction propensity of Tfrs with ns–GCBCs distracts the latter from receiving crucial survival signals from Tfhs. This led to a decreasing trend in the number and affinity of ns–ASCs(Fig. 4B,C, left panel). Of note, an indication of the same trend was appearing at the highest ratio in the case of the *Tfh-like* motility(Fig. 4A,B, right panel). When Tfrs obtained preferential access to the area rich in CCs with respect to Tfhs (*CC-like* motility), the selection of s-GCBCs was halted early during the GC reaction (Supplementary Material, Fig. S2B). The lack of s–GCBCs led to increased interaction between Tfrs and ns–GCBCs, diminishing the chance for ns–GCBCs to interact with Tfhs. This ultimately resulted in apoptosis of ns–GCBCs due to compromised collection of survival signals from Tfhs (Supplementary Material, Fig. S2B). Consequently, there was a decreasing trend in the number of GCBCs and ns–ASCs as the number of Tfrs were increased.

Thus, Tfr-induced s–GCBC apoptosis suppresses self-reactivity in GC efficiently even at low Tfr numbers, provided Tfrs can access the CCs-enriched area of the GC (Fig. 4D). However, under these assumptions, an optimally low number of Tfrs is necessary to avoid interference with the selection process of ns–GCBCs and guarantee a robust GC evolution. When Tfrs were dispersed within the LZ, their efficiency in inducing s–GCBC apoptosis and obstructing s–ASC differentiation was reduced. However, regardless of the spatial localization of Tfrs, the *Apoptosis* model indicates an optimal number of Tfrs to be beneficial in curbing self-reactivity in the GC while improving the affinity and numbers of ns–ASCs (Fig. 4D).

### 2.4 *In silico* Tfr-depletion reveals early Tfr-intervention in the context of Tfr-induced s–GCBC apoptosis

Considering the impact of Tfrs on ns–GCBCs, we explored the possibility of relieving the regulatory pressure of Tfrs on the selection process once the number of s–GCBCs decreases substantially. Our objective was to investigate whether this could facilitate the expansion and affinity maturation of ns– ASCs. Therefore, we simulated scenarios wherein Tfrs were depleted spanning three consecutive days, each starting at a specific stage of the GC reaction (Supplementary Material, Fig. S3). Consistent with the experimental conditions [18], the Tfr-depletion simulations were conducted with a Tfh:Tfr ratio of 2:1.

When Tfrs co-localized with CCs, depleting them shortly after the peak of the GC response (∼at day 7) resulted in a significant increase in the number of ASCs by ∼day 18 (Fig. 5A, right panel) without affecting their affinity (Fig. 5B, right panel). On the other hand, in the scenario where Tfrs were dispersed within the LZ resembling Tfh-like motility, their presence did not significantly impact the GC volume or the number of produced ASCs (Fig. 4A; Supplementary Material Fig. S4C). Consequently, depleting Tfrs after the first three days in the GC response did not lead to any significant changes (Fig. 5A, left panel). As an aside, a minor resurgence of s–GCBCs was observed in the case of Tfrs exhibiting *Tfh-like* motility, which was absent when Tfrs could access the CC-rich area of the GC (Supplementary Material, Fig. S3B,C, *Tfh-like* subpanels). These *in silico* experiments confirm that in the *Apoptosis* model, the suppressive function of Tfrs in controlling the expansion of s–ASCs is primarily exerted early in the GC response, at a time when the likelihood of generating and selecting s–GCBCs is higher.

**Figure 5:**
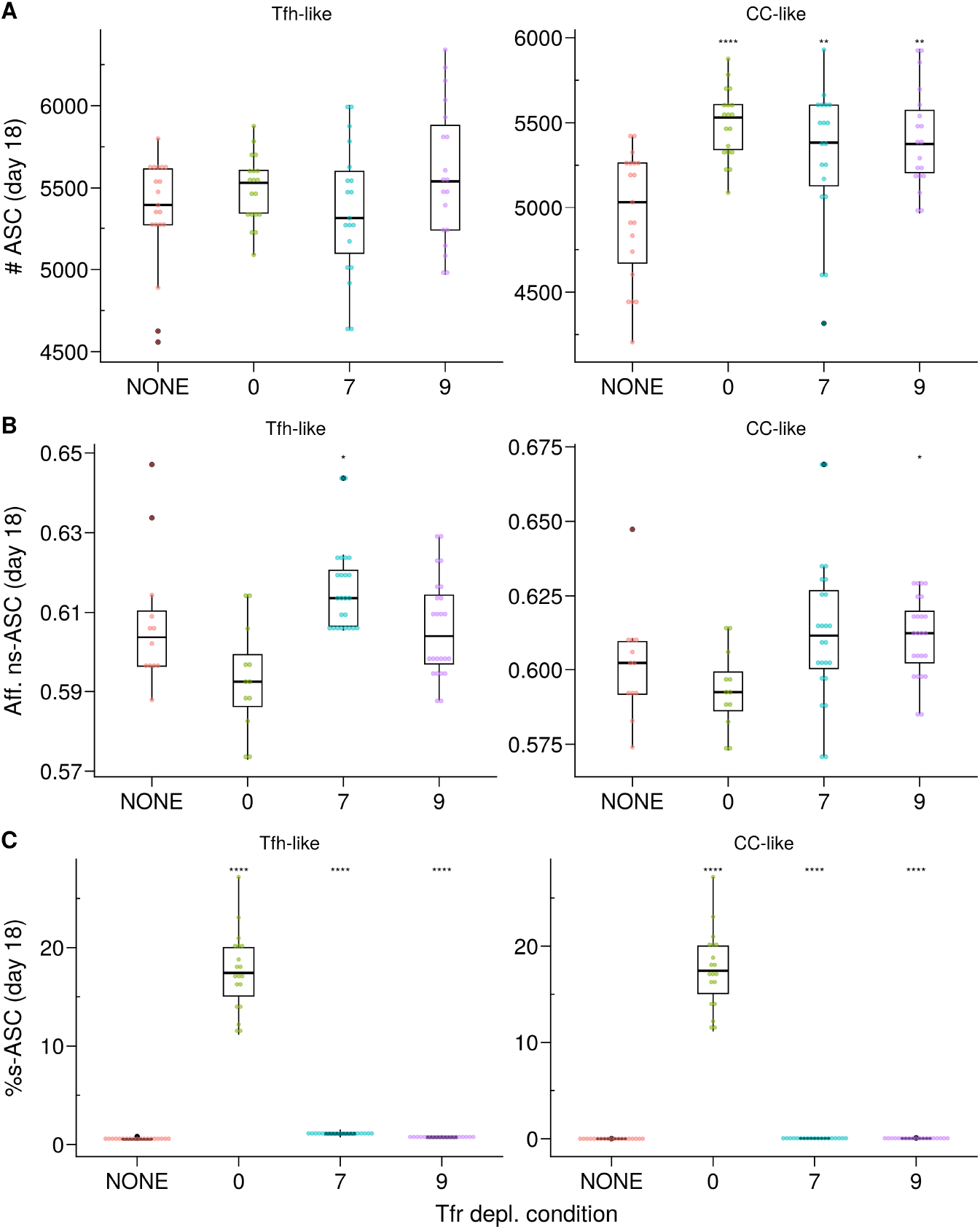
*In silico* Tfr depletion reveals early Tfr-intervention. **(A)** Number of ASCs; **(B)** Affinity of ns–ASCs; **(C)** s–ASCs percentage of total ASCs. Each column shows the results for the different motility types of Tfrs: *Tfh-like* (left coloumn), *CC-like* (right coloumn). Each colour denotes a different condition: no Tfr depletion (red), complete Tfr depletion pre-GC formation (green), partial Tfr depletion spanning 3 consecutive days starting at day 7 (cyan) or day 9 (purple). Each dot represents a different simulation. Significance of the findings is denoted with Wilcoxon signed rank tests against the *NONE* group. Statistics were performed on 20 simulations. The results were obtained at day 18 post-GC onset (∼day 21 post-immunization) using the *Apoptosis* model with *pSelf* = 0.04 and Tfh:Tfr = 2:1

To safeguard against autoimmunity, it is crucial to prevent the circulation of s–ASCs, irrespective of the events taking place within the GC. To address this, we analyzed how Tfrs can regulate B cell differentiation in the *SemiGate* and *SemiGate*.*38* models. This stage is critical as it determines the fate of the output cells as they prepare to exit the GC. Of note, the levels of engagement between Tfrs and GC B cells in both models were comparable regardless of the type of Tfr motility studied (Fig. 3B,C).

### 2.5 DZ recycling of s–GCPCs requires high Tfr numbers to contain s–ASCs and reduces ns–ASC population

In the context of the *SemiGate* model, Tfr interactions facilitated the differentiation of selected CCs, while s–GCPCs were enforced to undergo recycling in the DZ as a protective measure against the release of s–ASCs and the generation of self-reactive antibodies (Fig. 2C).

As previously observed, when *pSelf* = *pRed*, the proportion of B cells undergoing BCR re-editing to become non-self-reactive is minimal (Fig. 1C). Hence, s–GCBCs progressively accumulated within the GC over time (Supplementary Material, Fig. S4A), and we observed a minor increase of the GC volume, due to the increased number of recycling cells (Fig. 6A) and a corresponding reduction in the cumulative number of ASCs (Supplementary Material, Fig. S4C). Moreover, the accumulation of selected s–GCBCs resulted in a significantly decreased number of ns–ASCs (Fig. 6B), effectively preventing regular affinity maturation (Fig. 6C). This process entailed a higher number of Tfrs to cope with the increasing emerging wave of s–GCBCs.

**Figure 6:**
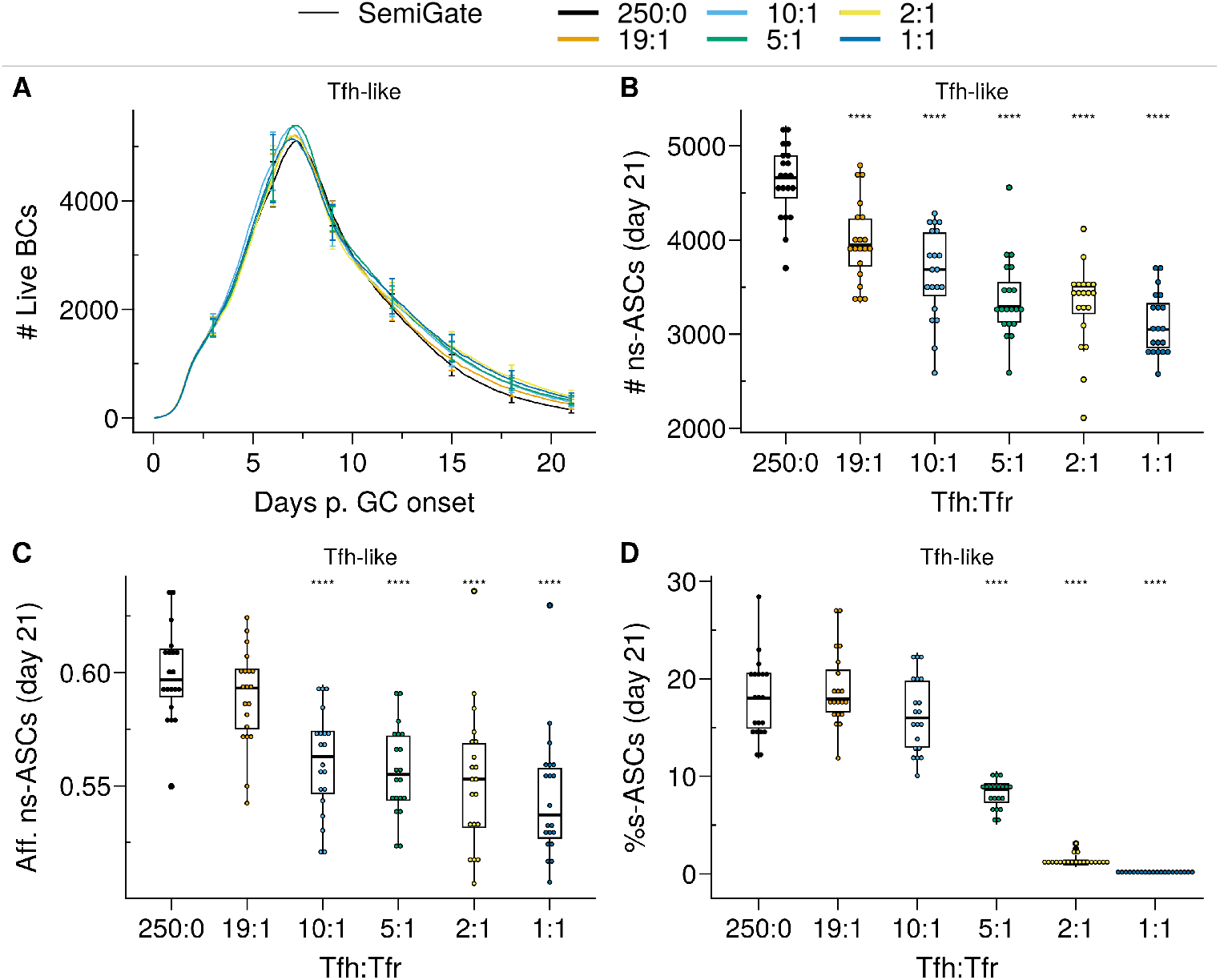
*SemiGate* model requires higher Tfr numbers to contain self-reactive antibody secreting cells (s–ASCs) and disrupts non-self-reactive ASCs (ns–ASCs). **(A)** Germinal centre B cell (GCBC) kinetics; **(B)** Number of ns–ASCs; **(C)** Affinity of ns–ASCs; **(D)** Percentage of s–ASCs in total ASCs. In **(B–D)** the analysis was performed at day 21 post-GC onset. Each colour depicts a different Tfh:Tfr ratio. In **(A)**, lines represent means and error bars show standard deviations at days 3,6,9,12,15,21. In **(B–D)**, each dot represents a different simulation and significance of the findings is denoted with Wilcoxon signed rank tests against the group with no Tfr (Tfh:Tfr = 250:0) to that with Tfrs. Statistics were performed on 20 simulations. The results were obtained with *pSelf* =0.04 for *Tfh-like* motility.

While increasing the number of Tfrs led to a significant reduction in s—ASCs (Fig. 6D), there was a gradual increase in the percentage of s–ASCs over time (Supplementary Material, Fig. S4B) due to the reinforcement of s–GCPC DZ recycling.

Thus, DZ recycling of s–GCPCs requires a high number of Tfrs to restrict the emergence of s–ASCs at the expense of the ns–ASC population and their affinity.

### 2.6 GCPCs engagement and s–ASC apoptosis by Tfrs suppressed self-reactivity without impacting ns–ASCs

In the *Semigate*.*38* model, Tfrs selectively interact with GCPCs, promoting their rapid differentiation and inducing apoptosis in s–ASCs post GC. The presence or absence of Tfrs had minimal impact on the overall evolution of the GC (Fig. 7A). Notably, Tfrs did not disrupt the GC selection process, thus having no significant impact on the numbers and affinity of ns–ASCs (Fig. 7B,C). Likewise, the dynamics of s–GCBCs remained comparable (Supplementary Material, Fig. S5A).

**Figure 7:**
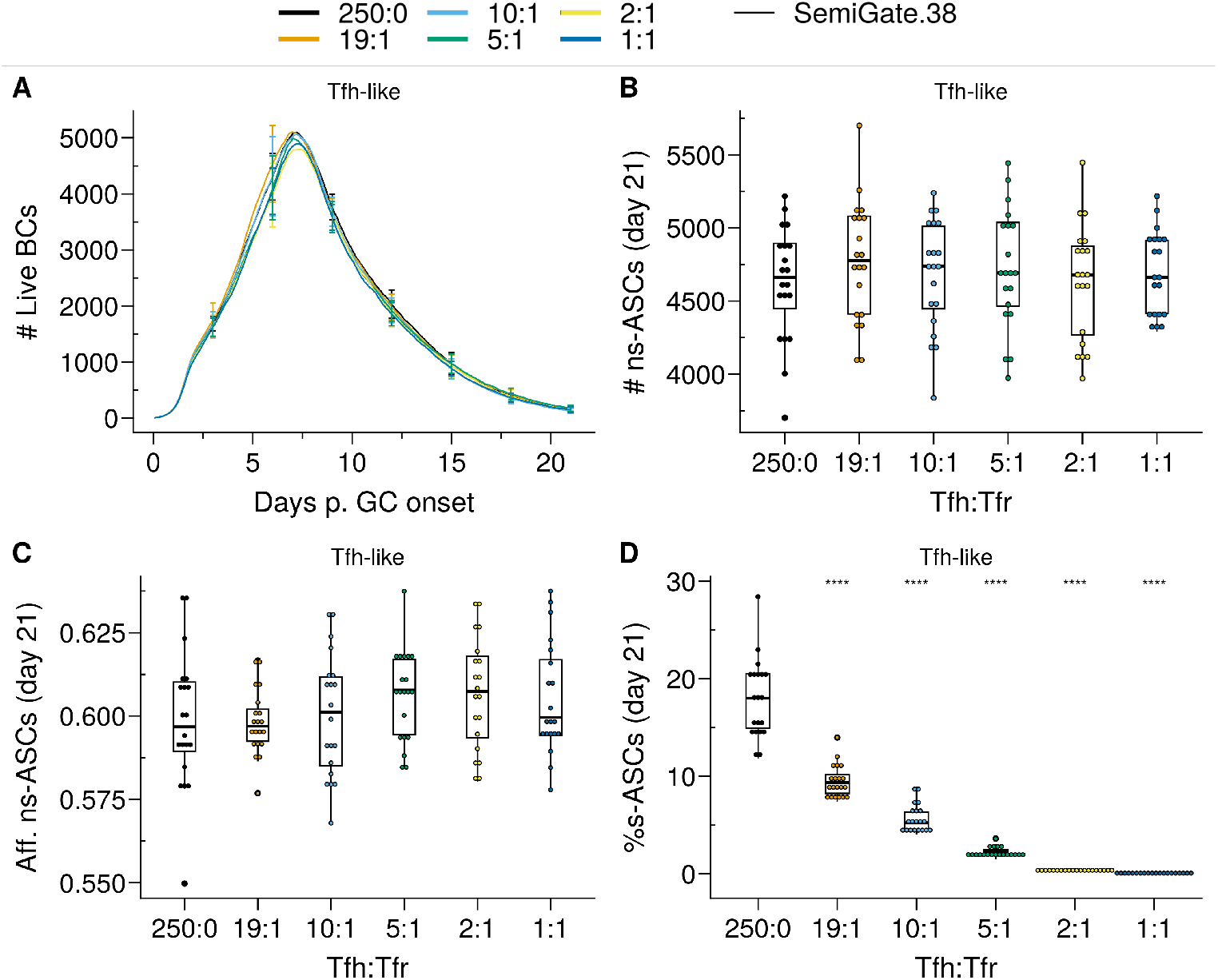
*SemiGate*.*38* model contains self-reactive antibody secreting cells (s–ASCs) in physiological Tfr numbers without impacting non-self-reactive ASCs (ns–ASCs). **(A)** Germinal centre B cell (GCBC) kinetics; **(B)** Number of ns–ASCs; **(C)** Affinity of ns–ASCs; **(D)** Percentage of s–ASCs in total ASCs. In **(B–D)** the analysis was performed at day 21 post-GC onset. Each colour depicts a different Tfh:Tfr ratio. In **(A)**, lines represent means and error bars show standard deviations at days 3,6,9,12,15,21. In **(B–D)**, each dot represents a different simulation and significance of the findings is denoted with Wilcoxon signed rank tests against the group with no Tfr (Tfh:Tfr = 250:0) to that with Tfrs. Statistics were performed on 20 simulations. The results were obtained with *pSelf* =0.04 and for *Tfh-like* motility of Tfrs.

As a result of Tfr-induced apoptosis in s–ASCs, there was a reduction in the total number of ASCs, which was primarily evident at higher Tfr numbers (Supplementary Material, Fig. S5C). Notably, even at low Tfr numbers (corresponding to high Tfh:Tfr), there was a significant reduction in the percentage of s–ASCs.

Taken together, the engagement of Tfrs with GCPCs and the subsequent induction of s–ASC apoptosis prove to be an efficient strategy in controlling the circulation of s–ASCs, even at low Tfr numbers, irrespective of their localization. Importantly, such a mechanism adversely affects neither the GC selection process nor the emergence of ns–ASCs.

Importantly, in the *Semigate* and *Semigate*.*38* models, Tfrs play a pivotal role in regulating the final differentiation of B cells as they prepare to exit the GC. As a consequence, unlike the *Apoptosis* model, these models require continuous monitoring and the presence of Tfrs within the GC. In both models, the population of s–GCBCs was maintained within the GC throughout the GC reaction (Supplementary Material, Fig. S4A,S5A). Consequently, the depletion of Tfrs led to the emergence of s–ASCs, with the extent of their emergence depending on the propensity of generating self-reactive B cell clones during SHM (Fig. 8A–D). Notably, both models exhibit a significant reduction in the percentage of GCPCs (Fig. 8D,E), similar to what was observed in experimental studies [19].

**Figure 8:**
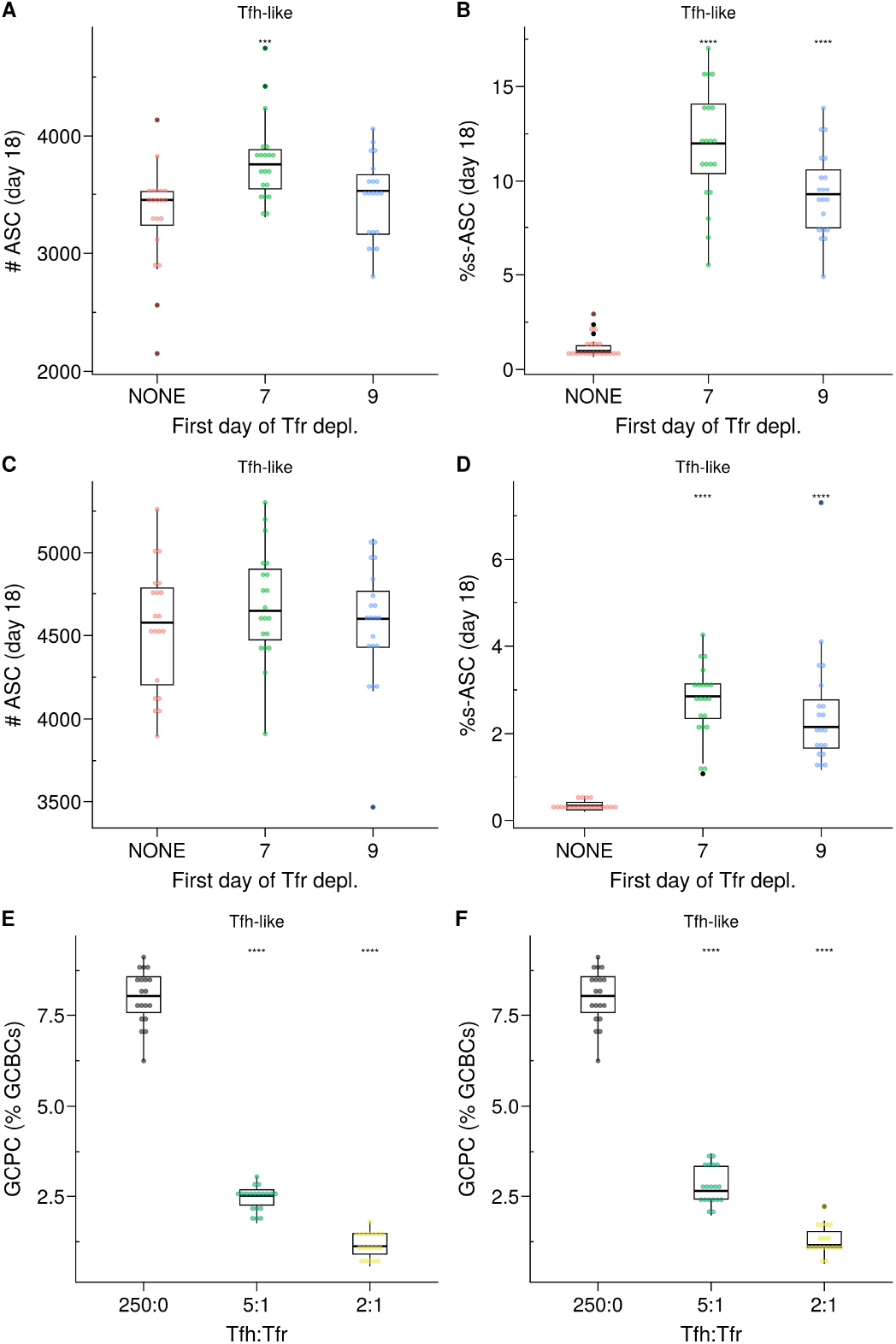
**(A–D)** *In silico* Tfr depletion experiment for the *SemiGate* **(A**,**B)** and for the *SemiGate*.*38* model **(C**,**D)** when Tfrs resemble *Tfh-like* motility. **(A**,**C)** Number of antibody secreting cells (ASCs); **(B**,**D))** Percentage of self-reactive ASCs (s–ASCs) in total ASCs. Each colour denotes a different condition: no Tfr depletion (red), partial Tfr depletion spanning 3 consecutive days starting at day 7 (green) or day 9 (blue). The analysis was performed at day 18 post-GC onset (∼day 21 post-immunization). Each dot represents a different simulation. Statistical significance is calculated with Wilcoxon signed rank tests against the *NONE* group. Statistics were performed on 20 simulations. **(E**,**F)** Germinal centre resident pre-plasma cells (GCPCs) in the *Semigate* and *Semigate*.*38* models, respectively. GCPCs are reported as a percentage of total germinal centre B cells (GCBCs) at day 14 post-GC onset (∼day 17 post-imm.) under different Tfh:Tfr ratios. Each dot represents a different simulation and statistical significance is calculated with Wilcoxon signed rank tests against the group with no Tfr (Tfh:Tfr = 250:0). Statistics were performed on 20 simulations. The results were obtained with *pSelf* = 0.04 and for *Tfh-like* motility of Tfrs.

## 3 Discussion

Despite extensive research elucidating the crucial role of Tfrs in regulating the development of self-reactive B cells in the context of the GC reaction, there still exist conflicting findings and an incomplete understanding of the underlying mechanism through which Tfrs exert their action. This raises the intriguing question how Tfrs effectively prevent self-reactivity given their relatively low numbers compared to their helper counterparts. In this study, we aimed to shed light on the suppressive function of Tfrs despite their limited abundance.

Our findings unveil the ability of Tfrs to exercise efficient control over self-reactivity by vigilantly monitoring the GC micro-environment and triggering apoptosis in s–GCBCs they interact with. Even under conditions with a higher likelihood of self-reactive B cells arising from SHM, Tfrs were able to impede the emergence of s–ASCs by inducing apoptosis in s–GCBCs early in the GC reaction (Fig. 4D). Notably, the presence of an optimal number of Tfrs facilitates the expansion and maturation of ns– GCBCs, thereby enhancing both the quantity and affinity of ns–ASCs (Fig. 4B,C).

When Tfrs had preferential access to the CC-rich region compared to Tfhs, even a lower number of Tfrs was sufficient to significantly suppress the emergence of s–ASCs. Interestingly, experiments identified a role of CCs in attracting the Tfrs by secretion of CCL3 to promote direct contacts between Tfrs and CCs [24]. In competitive settings, CCL3-KO BCs were increased with respect to their WT counterpart, indicating a suppressive role of Tfrs. The analysis on the role of CCL3 also showed that this chemokine might limit Tfr access to the DZ. Therefore, similar to Tfh [35], the location of Tfrs influences their efficiency in GC regulation.

In the *in silico* GC, as the number of Tfrs increased, their supportive role became detrimental to the GC evolution (Fig. 4A). Prominent engagement of Tfrs with ns–GCBCs reduced the opportunity of the latter to collect survival signals from Tfhs (Supplementary Material, Fig. S2B, *CC-like* motility panel). These findings indicate an intriguing role of Tfrs, which might act as distracting agents for ns– GCBCs during the selection process. Although the prominence of the distraction effect depends on the interaction time between Tfrs and ns–GCBCs, it indicates that the access of Tfrs into the GC must be controlled based on the potential for generating s–GCBCs, as suggested in experiments involving self-antigen immunization where an enhancement in Tfr numbers during the early stages of GC development resulted in GC shutdown GC [36]. On the other hand, insufficient access or inadequate presence of Tfrs in GCs during the later phase of the GC response may impede the timely shutdown of the GC. Notably, in autoimmune disorders, non-resolving GCs harbour distinct GC reactions initiated by B cell clones specific for auto-antigens and often exhibit a predominantly increased Tfh:Tfr ratio [37, 38, 39, 40, 41]. This suggests limited Tfr access to GCs or a potential inability to differentiate into Tfrs from Tfhs during the later stages of the GC response [25]. Of note, *in silico* GC shutdown is independent of any additional mechanism of differentiation of Tfhs to Tfrs.

Recent experiments revealed that halving the number of Tfrs had no significant impact on the GC volume or GC output at day 14 post-NP-KLH immunization [23]. It is worth noting that the Tfh:Tfr ratio in this experiment was within the higher range that we simulated (Tfh:Tfr = 10:1 or 20:1). Consistent with this result, increasing the Tfh:Tfr ratio from 10:1 to 19:1 yielded a similar GC evolution and number of output cells (Fig. 4).

Moreover, Tfrs were found to play a prominent role in regulating the size of the GC and ASCs, as well as the production of auto-antibodies during the initial phase of the GC response. Complete depletion of Tfrs before GC formation not only increased the number of GCBCs and GC output but also led to elevated levels of s–ASC differentiation (Fig. 5). Our *in silico* findings in the absence of Tfrs are consistent with these results, suggesting that Tfrs can indeed engage in cognate interactions with s–GCBCs and promote their apoptosis [36]. Depleting Tfrs at later stages of the GC reaction resulted in a greater resemblance to the GC phenotype observed in the absence of Tfr depletion (Fig. 5, day 9 of Tfr depletion).

As s–GCBCs were generated predominantly during the early days of the GC reaction, within the framework of the *Apoptosis* model, there was no significant difference in the proportion of auto-reactive BCRs among the live and apoptotic cells when analyzed after day 15 post-GC onset (Supplementary Material, Fig. S6A). However, the experimental data at day 10 to day 12 post-GC onset (∼day 14 post-immunization) revealed a similar proportion of auto-reactive BCRs in both the live and apoptotic compartments [32, 33], which disagreed with the findings from the *Apoptosis* model (Supplementary Material, Fig. S6A). In addition, this model was unable to reproduce the observed accumulation of GCPCs within the GC following Tfr-depletion (not shown). Furthermore, experimental findings suggesting suppression of auto-reactive cells in the post-GC phase indicate additional regulatory mechanisms after BC selection [33].

To explore how Tfrs may exert their regulatory actions during B cell differentiation, we examined two scenarios for containing s–ASCs. The *Semigate* model enforced the recycling of s–GCPCs into the DZ, whereas the *Semigate*.*38* model relied on post-GC apoptosis of s–ASCs. In the *Semigate* model, interaction with Tfrs resulted in prompt differentiation of selected CCs, whereas the *Semigate*.*38* model specifically engaged GCPCs with Tfrs, leading to their rapid differentiation (Fig. 2C,D).

In these models, the presence of Tfrs (Tfh:Tfr = 250:0) substantially reduced the accumulation of GCPCs within the GC (Fig. 8E,F), consistent with experimental findings [19]. Additionally, both models replicated the non-significant difference in the representation of auto-reactive BCRs between apoptotic and live compartments (Supplementary Material, Fig. S6B,C), as observed experimentally [32]. This can be attributed to Tfrs not directly interfering with the selection process of s–GCBCs. Consequently, in the *in silico* Tfr-depletion experiments with these models, there was a resurgence of s–GCBCs, as reflected in the proportion of s–ASCs (Fig. 8B,D; Supplementary Material, Fig. S4B, S5B). Nevertheless, these models demonstrated the ability to protect against self-reactive antibodies (Fig. 6C, 7C). However, this protection came at the cost of a reduced total quantity of ASCs (Supplementary Material, Fig. S4C, S5C), contradicting recent experimental observations [23].

Despite the aforementioned limitations of the *SemiGate* model, several experiments have demonstrated that Tfrs contribute to the promotion of Bcl6 expression and DZ phenotype [19, 22], as well as SHM of BCRs [42]. In the spirit of clonal redemption [29, 43, 44], we speculate that the interaction between s–GCPCs and Tfrs may enhance the DZ phenotype with a higher likelihood of mutation, allowing these cells to rescue their specificity for non-self. In our computational simulations, we indeed observed that this mechanism could redeem a greater number of self-reactive clones, thereby reducing self-reactivity (not shown). To obtain more conclusive insights and reconcile these experimental findings, a comprehensive analysis is needed to examine the impact of Tfr-GCBC interactions on B cell mutations.

In conclusion, we have developed an advanced and adaptable agent-based platform, which, to our knowledge, is the first of its kind to study the impact of Tfrs on various aspects of the GC reaction. Our results suggest two intriguing roles of Tfrs in the control of self-reactivity and the maturation of GC response. We unravel a possible reason for limiting the number of Tfrs, despite their essential role in controlling the emergence of s-ASCs. Our model suggests that non-physiological numbers of Tfrs interfere with the natural selection of ns-BCs, distracting them from collecting survival signals from Tfhs. Furthermore, our results hint that when the DZ phenotype of s-BCs is induced, Tfrs should promote mutation of these cells, thereby increasing the probability of BCR redemption.

## 4 Strengths and limitations of the study

Although we could not fully reconcile all the experimental evidence regarding how Tfrs suppress autoreactivity in the GC using a single model, this study underscores the importance of factors such as the spatial distribution of Tfrs, the relative abundance of s–GCBCs and ns–GCBCs, Tfrs and Tfhs, as well as the duration of interactions between B cells and Tfrs. These factors collectively determine the dynamics of Tfr engagement with B cells in GC, which in turn, dictate the dynamics and outcomes of the GC response, including the control of self-reactivity.

The presented study employs an abstract concept of self-reactivity, assuming that B cells undergo affinity maturation regardless of their specificity for self. As an aside, our computational models do not consider several established regulatory mechanisms employed by Tfrs, such as their impact on Tfhs and class switch recombination. Additionally, we did not implement the regulatory impact of cytokines and neuritin secreted by Tfrs explicitly. A systemic investigation integrating these regulatory factors with a dynamical model that coordinates the number, location and action of Tfrs may be imperative to reconcile the experimental findings.

## 5 Methods

### Germinal Centre Simulation Model

We describe the novel features included in the GC platform relevant for the current investigation. A complete description of the GC model can be found in the Supplementary Material.

### 5.1 B cell phenotypes

Three B cell phenotypes are distinguished: DZ B cells, LZ B cells, and Antibody Secreting Cells (ASCs). The different phenotypes characterise the cell properties and are not meant as localization within the GC zones. DZ B cells divide, mutate, and migrate. LZ B cells also migrate and undergo the different stages of the selection process. ASCs only migrate.

### 5.2 Self-reactive DZ B cells

Self-reactivity is represented as a property of the B cells. DZ B cells can acquire self-reactivity at each mutation (see Supplementary Material, *DZ B cells division*) with a fixed probability *pSelf*. The Self-reactivity property is inherited by the daughter cells. At each mutation, s–GCBCs have a probability of losing self-reactivity (*pRed*) equals to *pSelf*. Results are tested under different *pSelf* values.

### 5.3 Tfr cells

Tfrs are randomly distributed on the lattice and occupy a single node each. The simulation starts with a fixed number of Tfrs. Tfrs can move and interact with LZ B cells (CCs).

#### 5.3.1 Tfr motility

Tfrs migrate with an average speed of 10*µm/*min and repolarise every 1.7 minutes. Tfrs migrate differently depending on the motility-model.

- *CC-like*: Tfrs de- and resensitise for CXCL13 as described for LZ B cells in the *Chemotaxis* section of the Suppl;
- *Tfh-like*: Tfrs do random walk with a preferential directionality to the LZ, as described for Tfh in the *Chemotaxis* section.

### 5.4 Tfr Models

The state of the LZ B cells wich the Tfr can interact with and the outcome of the interaction depend on the *Tfr Model*. LZ B cells can be in the states unselected, FDC-contact, FDC-selected, TC-contact, TFR-contact, selected, apoptotic. We report the description of those states that were subjected to changes for the present investigation. A full description can be found in the Supplementary Material.

#### 5.4.1 FDC-Selected

LZ B cells in this state have acquired at least one antigen portion and can bind to either a Tfh or a Tfr. If Tfh and Tfr are neighbors of a B cell, the s–GCBC binds the Tfr, while the ns–GCBC binds the Tfh. If a LZ B cell meets a Tfh, it switches to the state *TC-contact*. In the *Apoptosis* model, if a LZ B cell meets a Tfr, it switches to the state *TFR-contact* (see *TFR-contact for the Apoptosis Model*).

#### 5.4.2 TFR-contact for the Apoptosis Model

In the *Apoptosis* model, LZ B cells interact with Tfr pre Tfh-selection. LZ s–GCBC remains bound to Tfr for 3 minutes [24, 25], and its accumulated Tfh-signaling time is reset to 0. After the binding time, s–GCBC detaches from the Tfr and returns to the state *FDC-selected* until the Tfh search time of 3 h is over. Further interactions with Tfhs or Tfrs are inhibited, and after the 3 h, they switch to the *apoptotic* state. LZ ns–GCBC remains immobile for 36 seconds. After the binding time, ns–BC detaches from the Tfr and returns to the state *FDC-selected*, and it continues to search and bind Tfhs or Tfrs until the Tfh search time of 3 h is over.

#### 5.4.3 Selected

Selected cells are desensitized to CXCL13 and perform a random walk. A probabilistic decision (0.2) is taken as to whether selected cells acquire a pre-PC state (GCPC) and differentiate to the ASCs phenotype after 12 hours (*differentiation delay*). The probability and time before differentiation were chosen to match the accumulation of CD138^+^ cells observed in the experiments in the absence of Tfr [19]. We refer to selected CCs which do not acquire a pre-PC state as CD138^*−*^ cells. CD138^*−*^ cells keep the LZ phenotype for 6 hours, then they differentiate to the DZ phenotype with a rate of 1/6 minutes. If a B cell recycles to the DZ phenotype, the delay in LZ B cell differentiation is counted as progression time in the cell cycle (corresponding to entering the cell cycle in the LZ). In the *SemiGate*.*38* and *SemiGate* models, only CD138^+^ cells and selected cells, respectively, can bind to Tfr. In these models, if a Tfr is neighbor of a B cell in the right state, the B cell switches to the state *TFR-contact* (see *TFR-contact for the Gating models*)

#### 5.4.4 TFR-contact for the *Semigate* and *Semigate*.*38* models

A selected B cell remains bound to the Tfr for 3 minutes and its *differentiation delay* is set to 0. After the binding time, the B cell detaches from the Tfr and returns to the state *selected* to complete its differentiation. In the *SemiGate* model, s–GCPCs return to the CD138^*−*^ cells phenotype. Therefore, they are forced to recycle to the DZ phenotype. In the *SemiGate*.*38* model, s–GCPCs are marked for death post GC (see *Antibody Secreting Cells*).

### 5.5 Antibody Secreting Cells

ASCs upregulate CXCR4, and leave the GC when they reach its boundary. In the *SemiGate*.*38* model, s–ASCs are deleted from the total number of ASCs after they leave the GC [33].

### 5.6 Tfr depletion experiment

Tfrs are depleted from the space at three consecutive days. Each day ∼40% of the present Tfrs are randomly deleted. In the main and supplementary figures, the results are reported for initial deletion at day 7 (∼peak of the GC) and day 9 (declining phase of the GC). The *NONE* case corresponds to simulations without Tfr depletion. The *0* case corresponds to simulations without Tfrs (Tfh:Tfr = 250:0).

## Author Contributions

*In silico* model construction, simulations and formal analysis: MS;

Investigation and discussion: MS, TM, AB, MMH;

Initial draft: MS, TM, AB;

Revision: TM, MS, MMH;

Supervision: MMH;

## Acknowledgments

We thank Prof. Carola Vinuesa for helpful suggestions and Dr. Sebastian C. Binder for reviewing the manuscript.

## Data Availability Statement

The original contributions presented in the study are included in the article and supplementary material. The codes developed and generated raw data analysed for this study can be obtained from the corresponding author upon request.

## Supplementary Material

A detailed description of the agent-based model of a germinal centre is provided in the Supplementary Material. In addition, supporting figures (Fig. S1–S6) can be found therein.

