## Supplementary Text and Figures for "Suppressive might of a few: T follicular regulatory cells impede auto-reactivity despite being outnumbered in the germinal centres"

† Corresponding authors

#### 1 Germinal Centre Simulation Model

The GC model used in the current work follows the description of [1] adapted to include novel features introduced since then in [2] and [3], as well as the self-reactive and the Tfr cells extension introduced here. Used acronyms are: DZ for dark zone, LZ for light zone, Tfh for T follicular helper cell, Tfr for T follicular regulatory cells, FDC for follicular dendritic cell, ns–GCBC and s–BC for non self-reactive and self reactive GC B cell, respectively, ns–ASC and s–ASC for non self-reactive and self reactive antibody secreting cell, respectively.

##### 1.1 Space representation

All reactions take place on a three-dimensional discretised space with a rectangular lattice with lattice constant of $\Delta x = 5\mu m$. Every lattice node can be occupied by a single cell only.

##### 1.2 Shape space for antibodies

Antibodies are represented on a four ($d = 4$) dimensional shape space [4]. The shape space is restricted to a size of 10 positions per dimension, thus, only considering antibodies with a minimum affinity to the antigen. The optimal clone $\varphi^*$ is positioned in the center of the shape space. A position on the shape space $\varphi$ is attributed to each B cell. The 1-Norm with respect to the optimal clone $\|\varphi - \varphi^*\|_1 = \sum_{i=1}^{d} |\varphi_i - \varphi_i^*|$, i.e., the minimum number of mutations required to reach the optimal clone, is used as a measure for the antigen binding probability. The binding probability is calculated from the Gaussian distribution with width $\Gamma = 2.8$ [5]:

$$b(\varphi, \varphi^*) = \exp\left( - \frac{\|\varphi - \varphi^*\|_1^2}{\Gamma^2} \right) \tag{1}$$

##### 1.3 B cell phenotypes

Three B cell phenotypes are distinguished: DZ B cells, LZ B cells, and Antibody Secreting Cells (ASCs). The different phenotypes characterise the cell properties and are not meant as localization within the GC zones. DZ B cells divide, mutate, and migrate. LZ B cells also migrate and undergo the different stages of the selection process. ASCs only migrate.

#### 1.4 Founder cells

The model starts from 250 Tfh, 200 FDCs, 300 stromal cells, a varying number of Tfr cells and no B cell. Tfh are randomly distributed on the lattice and occupy a single node each. Tfr position and movement is described in section *Tfr motility*. Stromal cells are restricted to the DZ (see section Chemokine distribution for their function). FDCs are restricted to the upper half of the reaction sphere, occupy one node by their soma and have 6 dendrites of $40\mu m$ length each. The presence of dendrites is represented as a lattice-node property and, thus, visible to B cells. The dendrites are treated as transparent for B cell or Tfh migration such that they do not inhibit cell motility.

#### 1.5 B cell influx rate

As B cell selection is not active during the first 3 days of the reaction (i.e., days 3-5 post immunisation), the first 3 days can be approximated as a B cell expansion phase. Clonality can be safely ignored, and it suffices to consider a single dividing cell type Bi, where i denotes the generation number of the B cells. The dynamics of expansion are then described by

$$\frac{dB_1}{dt} = s - pB_1$$

$$\frac{dB_i}{dt} = 2pB_{i-1} - pB_i \quad \text{for} \quad i > 1 \tag{2}$$

$$\frac{dB_{GC}}{dt} = 2pB_{i_{\max}},$$

where $s$ is the influx rate, $p$ the division rate, $i_{\max}$ the number of divisions per cell in the expansion phase, and $B_{GC}$ the resulting number of GC B cells that participate in the GC reaction. The number of initial divisions is estimated by the maximum number of divisions observed upon anti-DEC205-OVA treatment, i.e., $i_{\max} = 6$ [1, 6]. As the division time $\ln(2)/p$ is shorter than the expansion phase $T_{\text{expand}} = 3$ days, one may solve Eq.2 in steady state, yielding:

$$B6 = 2B5 = 4B4 = 8B3 = 16B2 = 32B1 = \frac{32s}{p} \tag{3}$$

Thus, the relevant ODE becomes

$$\frac{dB_{\text{GC}}}{dt} = 2pB_6 = 64s \quad \xrightarrow{\text{yields}} \quad B_{\text{GC}} = 64st \tag{4}$$

i.e., a linear growth in time proportional to the constant influx during expansion. Note that in the steady state approximation the influx rate becomes independent of the division rate $p$, which is an implication of the assumption of a fixed number of divisions per founder cell $i_{\max}$. With the side condition of getting 9000 cells at day 3, $B_{\text{GC}}(T_{\text{expand}} = 72\text{hr}) = 9000$, the influx rate is estimated to be

$$s = \frac{B_{\text{GC}}(T_{\text{expand}})}{64T_{\text{expand}}} \approx 2\frac{\text{cells}}{\text{hr}} \tag{5}$$

This corresponds to 144 B cells entering the GC in the first 3 days of expansion and building up the founder cell population of the GC reaction. Motivated by this estimation, in the model we assumed that B cells enter the GC reaction with a probability corresponding to a rate of 2 cells per hour for 96 hours. New B cells are randomly positioned on the lattice (exclusively on free nodes). The shape space position of each new B cell is randomly picked from a set of 100 shape space positions at a distance of 5 or 6 mutations to the antigen position.

#### 1.6 Antigen-presentation by FDCs

Each FDC is loaded with 3500 antigen portions distributed onto the lattice nodes occupied by FDC-soma or FDC-dendrite. One antigen portion corresponds to the number of antigen molecules taken up by a B cell upon successful contact with an FDC.

#### 1.7 DZ B cell division

The average cell cycle duration of 7 h of DZ B cells is varied for each B cell according to a Gaussian distribution. This is needed to get desynchronization of B cell division. The cell cycle is decomposed into four phases (G1, S, G2, M) in order to localize mitotic events if this is needed. Each founder B cell divides a number of times before differentiating to the LZ phenotype for the first time. Six divisions was the number of divisions found in response to the extreme stimulus with anti-DEC205-OVA [1, 6]. Each selected B cell divides a number of times determined by the interaction with Tfh (see below, LZ B cell selection). The parameters of the interaction with Tfh are tuned such that the mean number of divisions is in the range of two [7]. A division requires free space on one of the Moore neighbors of the dividing cell. Otherwise the division is postponed until a free Moore neighbor is available. At every division the encoded antibody can mutate with a probability of 0.5 [8, 9]. This corresponds to a shift in the shape space to a von Neumann neighbor in a random direction. Upon selection by Tfh the mutation probability is individually reduced from $m_{\max} = 0.5$ down to $m_{\min} = 0$ in an affinity-dependent way following

$$m(b) = m_{\max} - (m_{\max} - m_{\min})b = \frac{1-b}{2} \tag{6}$$

with b from Eq.1 [10]. Thus, after recycling DZ B cells can acquire reduced mutation probabilities. This mechanism is motivated by the observation that B cell receptor internalization enhances the activation of the kinase Akt [11] which, in turn, suppresses activation induced cytosine deaminase (AID) [12]. AID is required for somatic hypermutation, such that this provides an affinity dependent downregulation of the mutation frequency [13]. However, there is no formal proof of this mechanism. After the required number of divisions, the B cell differentiates with a rate of 1/6 min to the LZ phenotype.

#### 1.8 Self-reactive DZ B cells

Self-reactivity is represented as a property of the B cells. DZ B cells can acquire self-reactivity at each mutation with a fixed probability *pSelf*. The self-reactivity property is inherited by the daughter cells. At each mutation, s–BCs have a probability of losing self-reactivity (*pRed*) equals to *pSelf*. Results are tested under different *pSelf* values.

#### 1.9 LZ B cells

LZ BCs can be in the states unselected, FDC-contact, FDC-selected, TC-contact, TFR-contact, selected, apoptotic.

##### 1.9.1 Unselected

LZ B cells migrate and search for contact with FDCs loaded with antigen in order to collect antigen for 0.7 h. If an FDC soma or dendrite is present at the position of the BC, the BC attempts to establish contact to the FDC with probability b in Equation 1. Binding is affinity dependent and happens with the probability $b$ in Eq.1. If the available number of antigen portions at the specific FDC site drops below 20 the binding probability b is linearly reduced with the number of available portions. If successful, the B cell switches to the state *FDC-contact*; otherwise the B cell continues to migrate. Further binding-attempts are prohibited for 1.2 min. At the end of the antigen collection period, B cells switch to the state *FDC-selected*. If a LZ B cell fails to collect any antigen at this time it switches to the state *apoptotic*.

##### 1.9.2 FDC-Contact

LZ BCs remain immobile (bound) for 3 min [14] and then return to the state *unselected*. The counter for the number of successful antigen uptake events is increased by one and the FDC reduces its locally available antigen portions by one.

##### 1.9.3 FDC-Selected

LZ BCs in this state can bind to either a Tfh or a Tfr. If Tfh and Tfr are neighbors of a BC, the s–BC binds the Tfr cell, while the ns–BC binds the Tfh cell. If LZ BCs meet a Tfh they switch to the state *TC-contact*. In the *Apoptosis* model, if LZ BCs meet a Tfr they switch to the state *TFR-contact* (see *TFR-contact for the Apoptosis Model*).

##### 1.9.4 TC-contact

LZ BCs remain immobile for a time randomly sampled from a Gaussian distribution with mean 6 minutes and width 1.2 minutes. During this time, the bound Tfh cells, which may also be bound to other BCs, polarizes to the BC with highest number of successful antigen uptakes and polarization is recalculated in every time step. If more than one BC shares the same highest pMHC presentation, polarization is chosen randomly among those BCs. Only this B cell receives Tfh signals and accumulate those. After the binding time, the B cell detaches and returns to the state *FDC-selected*. It continues to search and bind Tfh cells until the Tfh search time of 3 h is over. Then, it switches to the state *apoptotic* if the accumulated Tfh-signaling time remained below 18 min. Otherwise it switches to the state *selected*.

##### 1.9.5 TFR-contact for the Apoptosis Model

In the *Apoptosis* model, LZ BCs interact with Tfr pre-Tfh-selection. LZ s–BC remains bound to Tfr for 3 minutes [15, 16], and its accumulated Tfh-signal is reset to 0. After the binding time, s–BC detaches from the Tfr and returns to the state *FDC-selected* until the Tfh search time of 3 h is over. Further interactions with Tfhs or Tfrs are inhibited, and after the 3 h, they switch to the *apoptotic* state. LZ ns–BC remains immobile for 36 seconds. After the binding time, ns–BC detaches from the Tfr and returns to the state *FDC-selected*, and it continues to search and bind Tfhs or Tfrs until the Tfh search time of 3 h is over.

##### 1.9.6 Selected

Selected cells are desensitized to CXCL13 and perform a random walk. A probabilistic decision (0.2) is taken as to whether selected cells upregulate CD138 (GCPC) and differentiate to the ASC phenotype after 12 hours (*differentiation delay*). The probability and the *differentiation delay* were chosen to match the accumulation of $CD138^+$ cells observed in the experiments in the absence of Tfrs [17]. $CD138^-$ cells keep the LZ phenotype for 6 hours. Then, they differentiate to the DZ phenotype with a rate of 1/6 minutes. If a BC recycles to the DZ phenotype, the *differentiation delay* is counted as progression time in the cell cycle (corresponding to entering the cell cycle in the LZ). In the *SemiGate* and *SemiGate.38* models, only selected cells and only $CD138^+$ cells, respectively, can bind to Tfrs. If a Tfr cell is a neighbour of a BC, BC switches to the state *TFR-contact* (see *TFR-contact for the Gating models*).

Recycling BCs memorize the amount of integrated Tfh-signal as well as the cell cycle phase they have reached by the selection time. The number of divisions $P(C)$ the recycled BCs will do is derived from the amount of integrated Tfh cell signal $C$,

$$P(C) = P_{\rm min} + (P_{\rm max} - P_{\rm min}) \frac{C^{n_{\rm P}}}{C^{n_{\rm P}} + K_{\rm P}^{n_{\rm P}}} \tag{7}$$

The minimum number of division is set to one $P_{\rm min} = 1$ in order to avoid recycling events without further division. It is limited by 6 divisions in the best case, which is motivated by anti-DEC205-OVA experiments in which DEC205+/+ BCs received abundant antigen which increased pMHC presentation to a maximum [6]. The population dynamics *in vivo* and *in silico* only matched when the number of divisions was increased to six in the simulation [1] suggesting that the strongest possible pMHC presentation to Tfh cells induces six divisions $P_{\rm max} = 6$. The Hill-coefficient was $n_{\rm p} = 1.2$.

The half value $K_{\rm P}$ remained to be determined, which denotes the amount of signal collected by B cells at which the number of divisions becomes half maximal. It was fixed by the condition that for the mean level of $C = C_0$ the number of divisions becomes two, as observed in [7]:

$$K_{\rm P} \approx C_0 \left( \frac{P_{\max} - P_{\min}}{P_0 - P_{\min}} - 1 \right)^{1/n_{\rm P}}. \tag{8}$$

with $C_0 = 0.45$ and $P_0 = 2$.

###### 1.9.7 TFR-contact for the Gating models

Selected BC remains bound to the Tfr cell for 3 minutes, and its *differentiation delay* is set to 0. In the *SemiGate* model, s–GCPCs returns as $\text{CD138}^-$ cells phenotype. Therefore, they are forced to recycle to the DZ phenotype. In the *SemiGate.38* model, s–GCPCs are marked for death post-GC. After the binding time, BC detaches from the Tfr and returns to the state *selected* to complete its differentiation.

###### 1.9.8 Apoptosis

LZ BCs remain on the lattice for 6 hours before they are deleted.

##### 1.10 Antibody Secreting Cells

ASCs upregulate CXCR4 and leave the GC when they reach the GC boundary. In the *SemiGate.38* model, s–ASCs are deleted from the total number of ASCs after they leave the GC [18].

##### 1.11 Chemokine distribution

Two chemokines CXCL12 and CXCL13 are considered. CXCL13 is produced by FDCs in the LZ with 10nMol per hour and FDC while CXCL12 is produced by stromal cells in the DZ with 400nMol per hour and stromal cell. As both cell types are assumed to be immobile, chemokine distributions were pre-calculated once and the resulting steady state distributions were used in all simulations.

##### 1.12 Chemotaxis

DZ and LZ BCs regulate their sensitivity to CXCL13 and CXCL12, respectively. This is true in all BC states unless stated otherwise. All BCs move with a target speed of 7.5$\mu$m/min. This leads to a slightly lower observable average speed of $\sim 6\mu$m/min. BCs have a polarity vector that determines their preferential direction of migration. The polarity vector $\vec{p}$ is reset every 1.5 minutes into a new direction using the chemokine distribution $c$ as

$$\vec{p} = \vec{p}_{\rm rand} + \frac{\alpha}{1 + \exp\left\{ \kappa \left( K_{1/2} - \Delta x \left| \vec{\nabla} c \right| \right) \right\}} \frac{\vec{\nabla} c}{\left| \vec{\nabla} c \right|}, \tag{9}$$

where $\vec{p}_{\rm rand}$ is a random polarity vector and the turning angle is sampled from the measured turning angle distribution ([19], Figure S1B). $\alpha = 3$ determines the relative weight of the chemotaxis and random walk, $K_{1/2} = 2 \cdot 10^{11}$Mol determines the gradient of half maximum chemotaxis weight, and $\kappa = 10^{10}$/Mol determines the steepness of the weight increase.

BCs de- and re-sensitize for their respective chemokine depending on the local chemokine concentration: The desensitization threshold is set to 4.5 nMol and 0.08 nMol for CXCL12 and CXCL13, respectively, which avoids cell clustering in the center of the zones. The resensitization threshold is set at 2/3 and 3/4 of the desensitization threshold for CXCL12 and CXCL13, respectively. BCs can only migrate if the target node is free. If occupied and the neighbor cell is to migrate in the opposite direction (negative scalar product of the polarity vectors) both cells are exchanged with a probability of 0.5. This exchange algorithm avoids lattice artifacts leading to cell clusters. Tfh cells do random walk with a preferential directionality to the LZ: the polarity vector $\vec{p}$ of Tfh cells is determined from a mixture of random walk $\vec{r}$ and the direction of the LZ $\vec{n}$ by

$$\vec{p} = (1 - \alpha')\vec{r} + \alpha'\vec{n}, \tag{10}$$

where $\alpha' = 0.1$ is the weight of chemotaxis. This weight leads to a dominance of random walk with a tendency to accumulate in the LZ as found in experiment. Tfh cells migrate with an average speed of $10\mu m/\text{min}$ and repolarize every 1.7 minutes [20]. ASCs motility is derived from plasma cell motility data to $3\mu m/\text{min}$ with a persistence time of 0.75 minutes [19].

#### 1.13 Tfr Motility

Tfr cells migrate with an average speed of $10\mu m/\text{min}$ and repolarise every 1.7 minutes. Tfr cells migrate differently depending on the motility-model.

##### 1.13.1 CC-like

Tfr are randomly distributed on the lattice and occupy a single node each. Tfr cells de- and resensitise for CXCL13 as described for LZ B cells in the *Chemotaxis* section.

##### 1.13.2 Tfh-like

Tfr are randomly distributed on the lattice and occupy a single node each. Tfr cells do random walk with a preferential directionality to the LZ, as described for Tfh in the *Chemotaxis* section.

### 2 Supplementary Figures

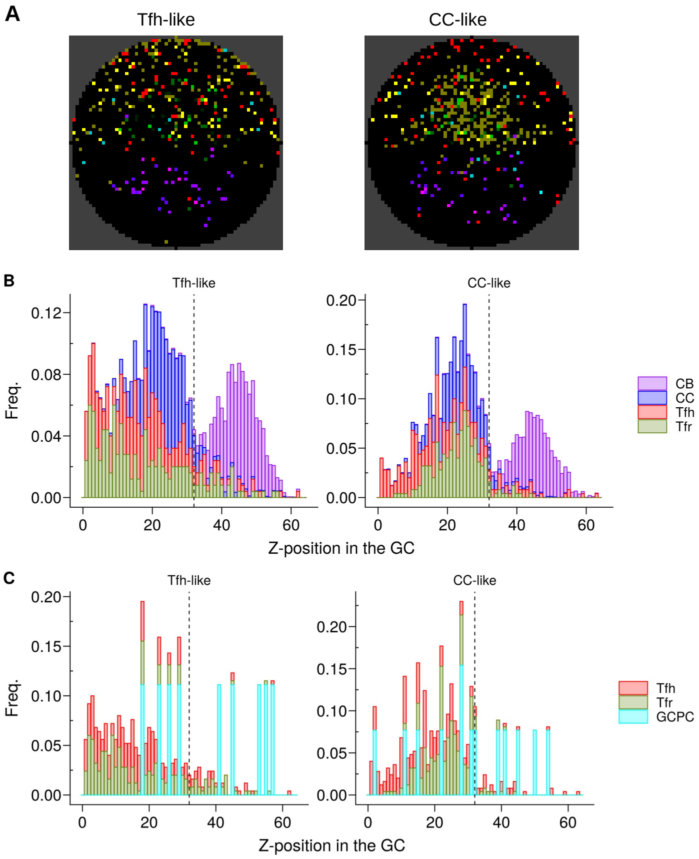


Figure 1: Tfr distribution in the GC: **(A)** Representative *in silico* GC sections at day 3-4 post-GC onset; LZ and DZ are the upper and lower half of the circle, respectively; different colours represent different types of cells: Tfhs (red), Tfrs (green), FDCs stroma (yellow), CBs (purple), CCs (green); **(B,C)** Representative frequency of cells at each Z-positions in the GC at day 10 post-GC onset. Colours indicate different types of cells and vertical dotted black line defines LZ (left)–DZ (right) limit. **(A–C)** Results shown for *Tfh-like* motility (left coloumn) and *CC-like* motility (right coloumn) of Tfrs under Tfh:Tfr = 2:1.

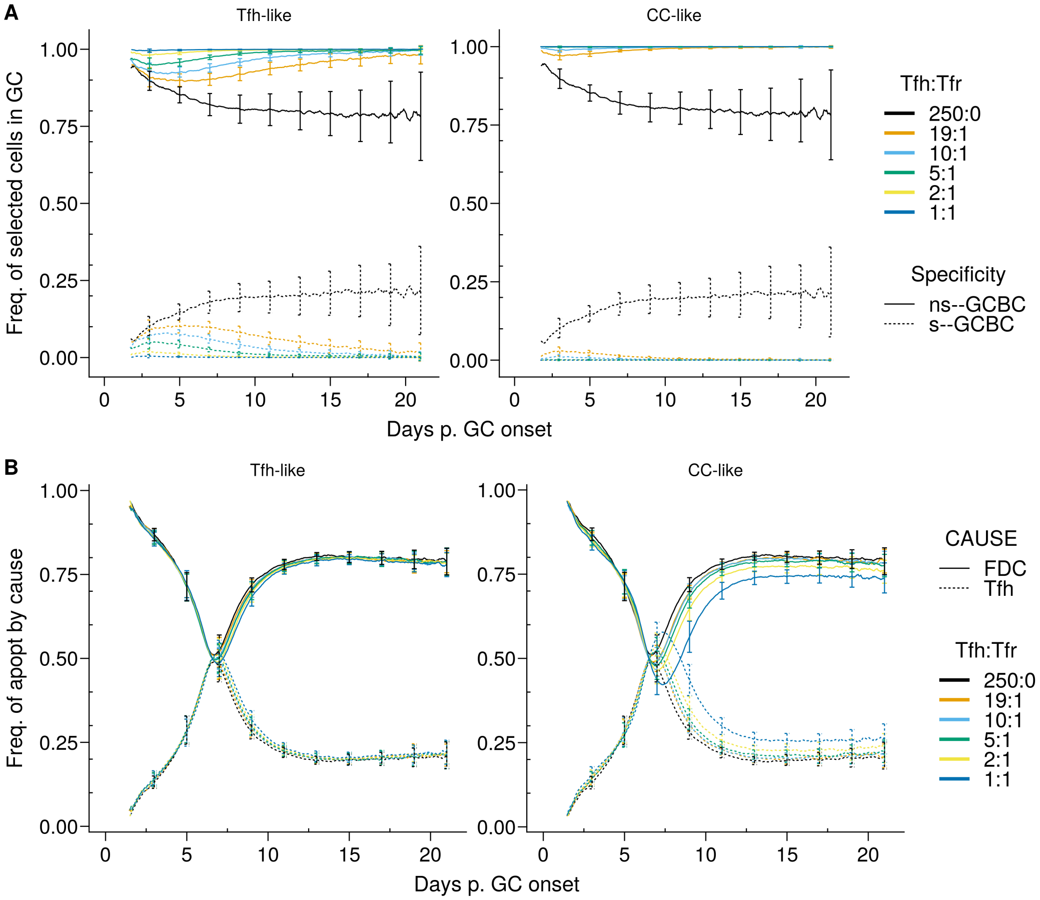


Figure 2: **(A)** Time-evolution of the selection frequency of cells with different specificities, self (s–GCBC) non-self (ns–GCBC). **(B)** Time-evolution of frequency of the cause of apoptosis, failure of antigen acquisition (FDC), lack of Tfh help (TFH). Each coloumn shows the results for the different motility types: *Tfh-like* (left coloumn), *CC-like* (right coloumn). Each colour depicts a different Tfh:Tfr ratio. Lines and error bars are means and standard deviation, respectively. SD at days 3,6,9,12,15,21. Statistics performed on 96 simulations. The results were obtained with *pSelf*=0.04.

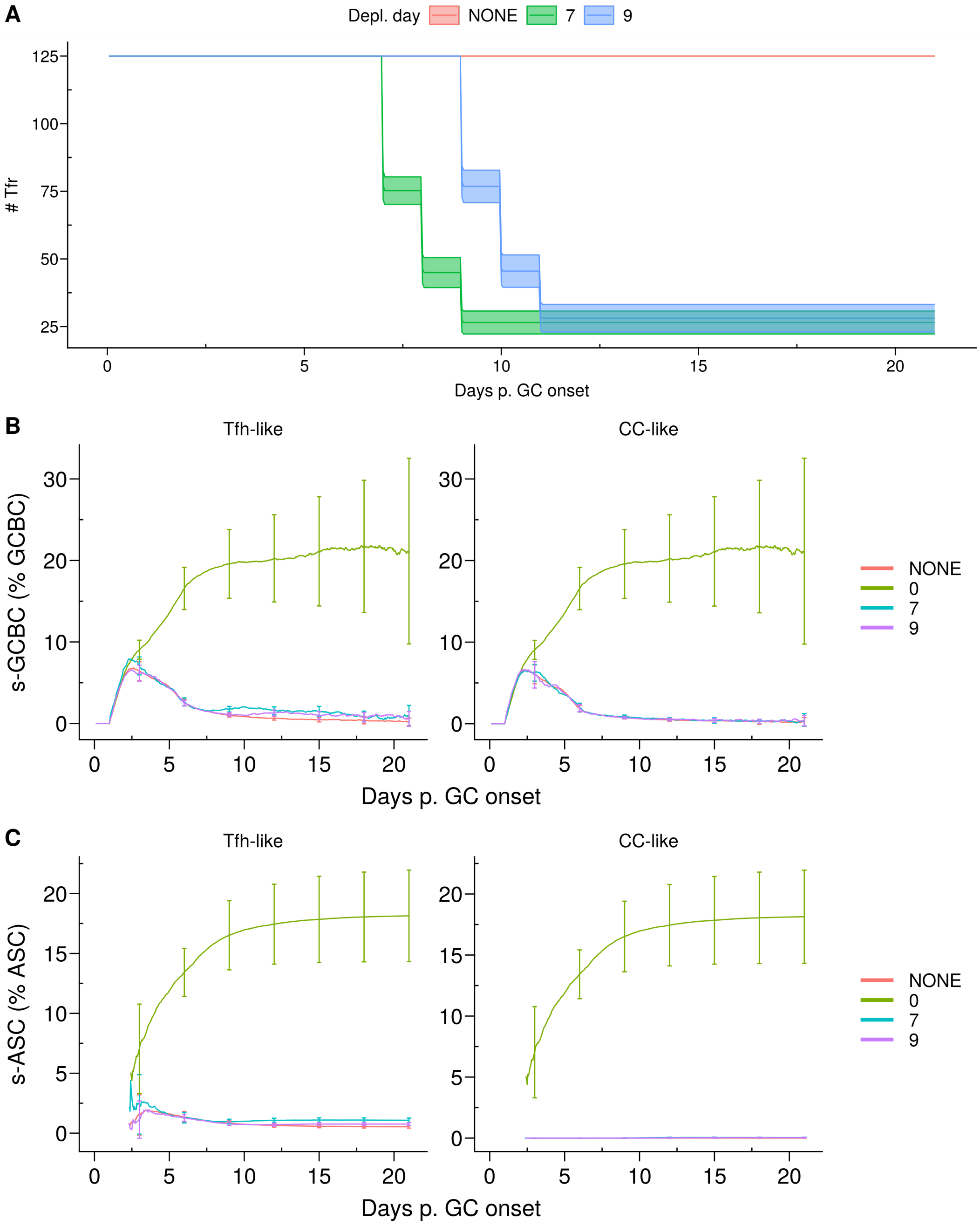


Figure 3: *In silico* Tfr depletion experiment. **(A)** Tfr number post *in silico* Tfr depletion experiment. Tfr were depleted at three consecutive days, reaching ~1/5 of the initial value. **(B)** Kinetics of s–GCBCs as percentage of total GCBCs; **(C)** Kinetics of s–ASCs as percentage of total ASCs. Each coloumn shows the results for the different motility types: *Tfh-like* (left coloumn), *CC-like* (right coloumn). The experiment was performed starting at different days post-GC onset (Depl. day) and each colour is a different day at which depletion was initiated. Lines and shaded areas **(A)** or error bars **(B–C)** are mean and standard deviations, respectively, of 96 simulations. The results were obtained using the *Apoptosis* model, with *pSelf* = 0.04 and Tfh:Tfr = 2:1.

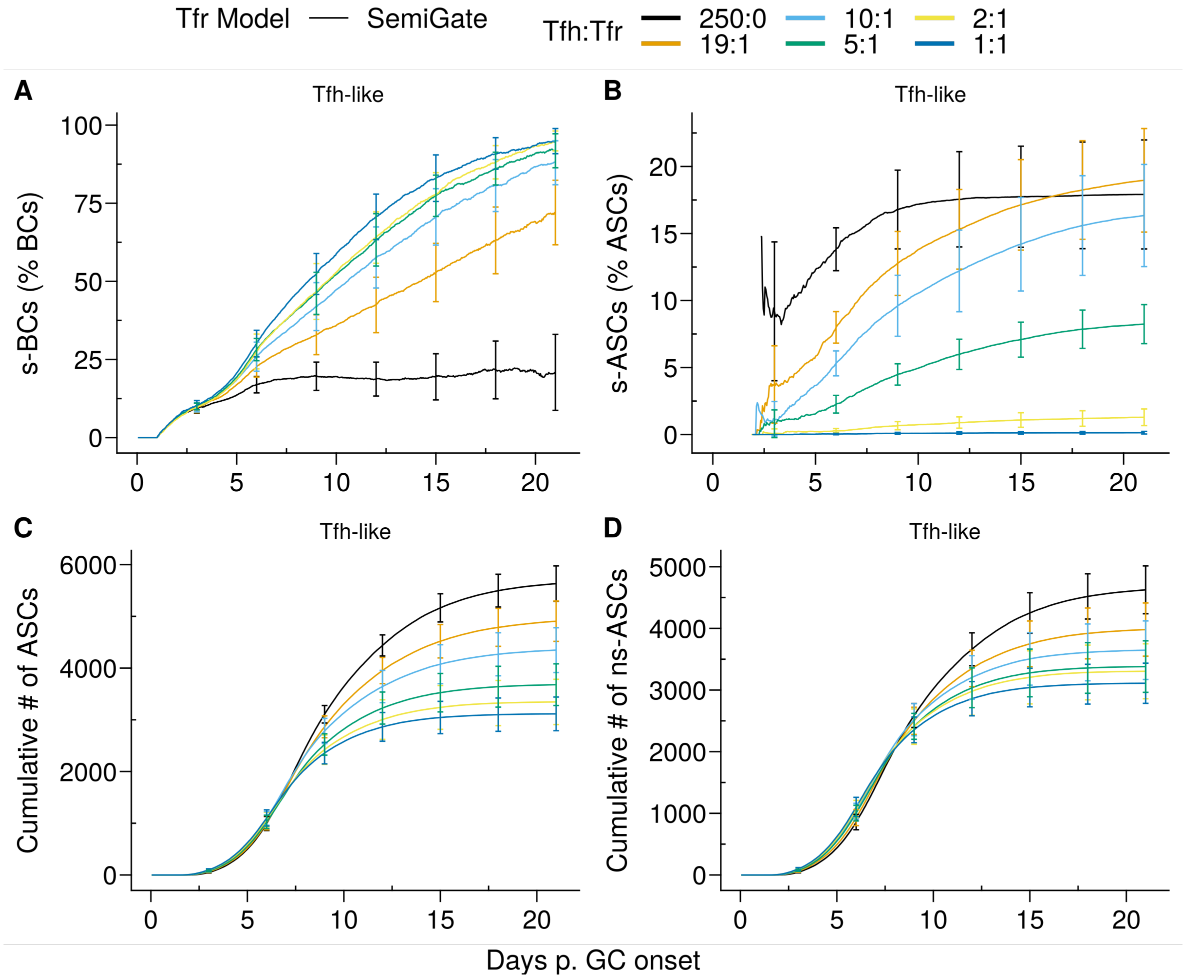


Figure 4: *SemiGate* model. Kinetics of **(A)** s–GCBCs as percentage of total GCBCs, **(B)** s–ASCs as percentage of total ASCs, **(C)** cumulative number of ASCs and **(D)** cumulative number of ns–ASCs. Each colour depicts a different Tfh:Tfr ratio. In **(A–D)**, lines and error bars are means and standard deviations, respectively. SD at days 3,6,9,12,15,21.

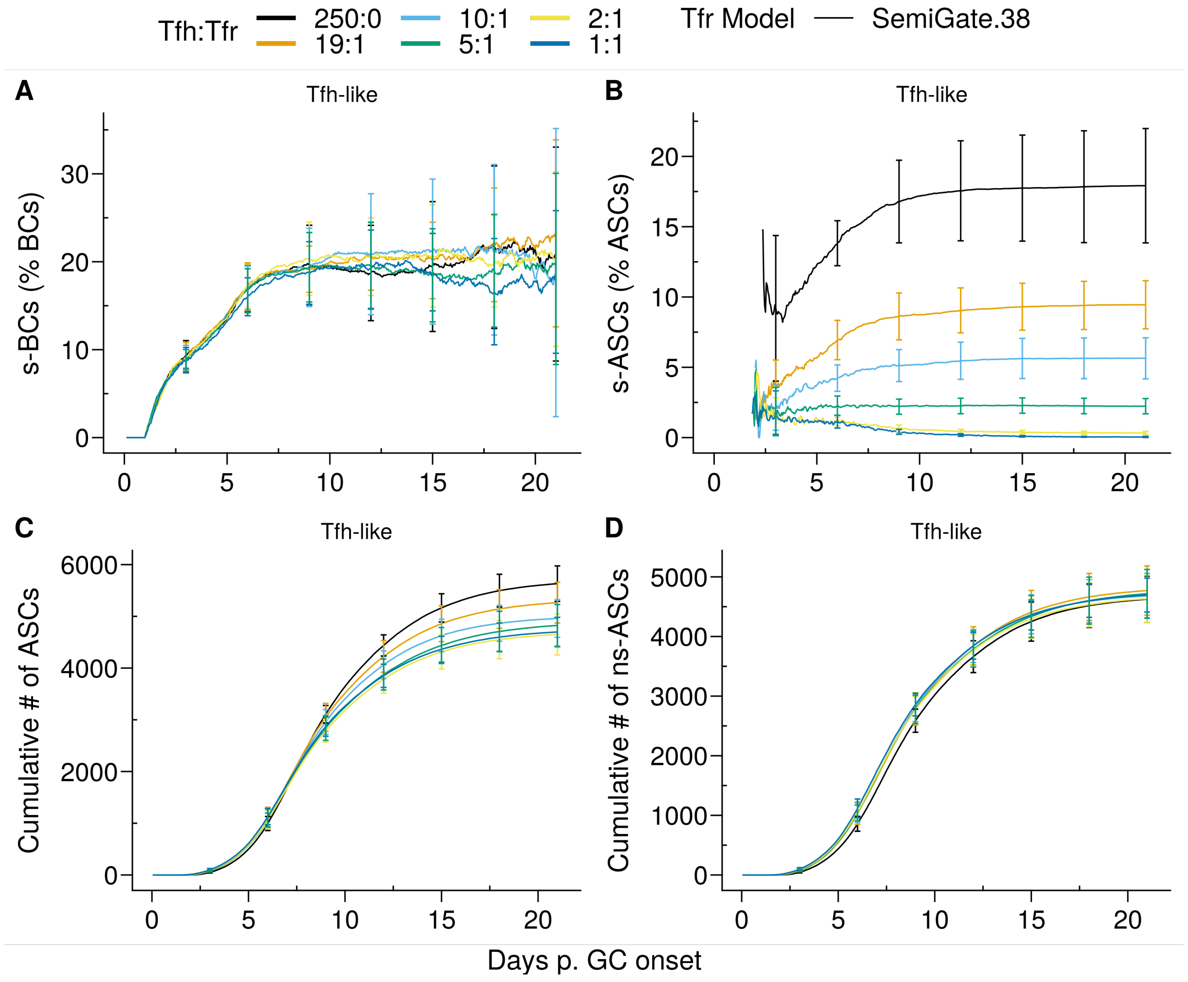


Figure 5: *SemiGate.38* model. Kinetics of **(A)** s–GCBCs as percentage of total GCBCs, **(B)** s–ASCs as percentage of total ASCs, **(C)** cumulative number of ASCs and **(D)** cumulative number of ns–ASCs. Each colour depicts a different Tfh:Tfr ratio. In **(A–D)**, lines and error bars are mean and standard deviation, respectively. SD at days 3,6,9,12,15,21.

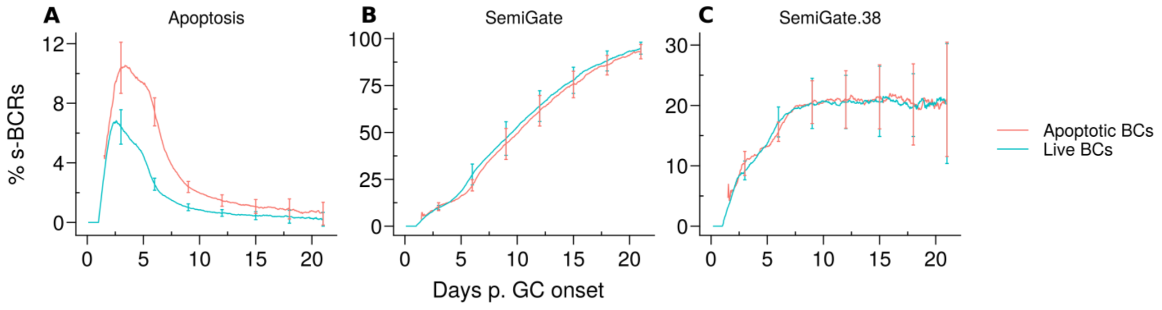


Figure 6: BCR specificity. Kinetics of the s–GCBCs as a % of the total GCBCs in the compartment of live BCs (cyan) and apoptotic BCs (red) for **(A)** the *Apoptosis* model, **(B)** the *SemiGate* model and **(C)** the *SemiGate.38* model. Lines and error bars are mean and standard deviation, respectively, performed on 20 simulations. The results were obtained with $pSelf = 0.04$, under for *Tfh-like* motility of Tfrs and Tfh:Tfr = 2:1.
